# Soma-centered control of synaptic autophagy by Rab39-regulated anterograde trafficking of Atg9

**DOI:** 10.1101/2024.11.21.624639

**Authors:** Ayse Kilic, Dirk Vandekerkhove, Sabine Kuenen, Jef Swerts, Esther Muñoz Pedrazo, Carles Calatayud Aristoy, Abril Escamilla Ayala, Nikky Corthout, Pablo Hernández Varas, Stephane Plaisance, Eliana Nachman, Valerie Uytterhoeven, Patrik Verstreken

## Abstract

Presynaptic terminals can be located far from the neuronal cell body and are thought to independently regulate protein and organelle turnover. In this work, we report a soma-centered mechanism that regulates autophagy-driven protein turnover at distant presynaptic terminals in *Drosophila*. We show that this system is regulated by Rab39, whose human homolog is mutated in Parkinson’s disease. Although Rab39 is localized in the soma, its loss of function causes increased autophagy at presynaptic terminals, resulting in faster synaptic protein turnover and neurodegeneration. Using a large-scale unbiased genetic modifier screen, we identified genes encoding cytoskeletal and axonal organizing proteins, including *Shortstop* (*Shot*), as suppressors of synaptic autophagy. We demonstrate that Rab39 controls Shot-and Unc104/KIF1a-mediated transport of autophagy-related Atg9 vesicles to synapses. Under starvation conditions, Rab39 in the soma shifts its localization from endosomes to lysosomes, thereby controlling the availability of Atg9 vesicles for trafficking to synapses. Our findings indicate that Rab39-mediated trafficking in the soma orchestrates a cross-compartmental mechanism that regulates the abundance of autophagy at synapses.

## Introduction

Synaptic autophagy is essential for maintaining neuronal health by ensuring the clearance and recycling of dysfunctional synaptic components (Soukup et al. 2016; Bademosi et al. 2023; Vijayan and Verstreken 2017; Birdsall and Waites 2019). This process is likely crucial for coping with the high metabolic activity and required for the functional integrity of synapses and effective neuronal communication. Given that synapses are often located far from the cell body, they appear to independently regulate this process. It remains unclear whether there is also cell body oversight in the regulation of synaptic autophagy.

Several proteins mutated in Parkinson’s disease (PD) play key roles in regulating autophagy at synapses (Grosso Jasutkar and Yamamoto 2023; Nachman and Verstreken 2022). Leucine-rich repeat kinase 2 (LRRK2)/Lrrk-dependent phosphorylation of the synaptic protein EndophilinA (EndoA) (Matta et al. 2012) drives the process in a starvation and activity dependent manner, and PD pathogenic mutations in LRRK2 and EndoA disrupt autophagy at presynaptic terminals (Bademosi et al. 2023; Soukup et al. 2016; D. Chang et al. 2017). Similarly, EndoA binds to several PD-linked proteins (Watanabe et al. 2018), including Parkin and Synaptojanin-1, which are also both causally mutated in PD. Synaptojanin-1, like EndoA, regulates vesicle trafficking and autophagy specifically at synapses and not in neuronal cell bodies (Cao et al. 2014; Vanhauwaert et al. 2017; Song et al. 2023; Yang et al. 2022). Interestingly, in PD, neurodegeneration is believed to begin with subtle synaptic defects before leading to overt neuronal death. (Schirinzi et al. 2016) and deregulation of synaptic autophagy, including the pathogenic mutation in EndoA, causes neurodegeneration *in vivo* (Bademosi et al. 2023; Soukup et al. 2016; Grosso Jasutkar and Yamamoto 2023). Interestingly, both excessive and insufficient synaptic autophagy have the same detrimental outcome, highlighting the importance of maintaining a finely tuned balance of this process to ensure neuronal health.

Human genetic evidence links vesicle trafficking defects to PD. Rab GTPases are crucial regulators of cellular trafficking events (Schimmöller, Simon, and Pfeffer 1998) and Genome-Wide Association Studies (GWAS) have identified variants at the *RAB29* locus that increase the risk of PD (Kia et al. 2021; Nalls et al. 2019; Tan et al. 2024). Additionally, a rare mutation in another Rab-GTPase, *RAB32*, also increases the risk for PD (Gustavsson et al. 2024; Hop et al. 2024) and familial mutations implicate the loss of function of *RAB39B*, as causative to the disease (Gao, Wilson, Salce, et al. 2020; Jacobson et al. 2024). Moreover, LRRK2 has been shown to mediate the phosphorylation of various Rabs, including Rab3, Rab8, Rab10, Rab12, and possibly Rab29 (Steger et al. 2016; Fujimoto et al. 2018; Liu et al. 2018; Purlyte et al. 2018; Dou, Aiken, and Holzbaur 2024). While the connection of PD to Rab GTPases is evident, their potential relation to synaptic EndoA-dependent-autophagy remains less well understood.

Here, we demonstrate that similarly to a *Rab39* knock-out, PD-associated mutations in *Rab39B* result in increased synaptic autophagy, akin to the effects observed with the *LRRK2^G2019S^* or phosphomimetic *endoA^S75D^*mutants (Soukup et al. 2016). To investigate the regulation of this process, we utilized the *endoA^S75D^* mutant in a large-scale unbiased genetic modifier screen in *Drosophila*. This approach uncovered key regulators of synaptic autophagy, indicating that the loss of function in genes involved in cytoskeletal organization and axonal transport suppress the increased autophagy in both *endoA^S75D^* and *Rab39* mutants. While EndoA localizes directly at synapses, Rab39 is primarily restricted to the neuronal cell body (Chan et al. 2011; Gao et al. 2020). Strikingly, we found that Rab39 within the soma plays a critical role in inhibiting autophagy at the synapse by controlling the trafficking of Atg9-positive vesicles. Disruption of this Rab39-dependent trafficking pathway results in increased Atg9 transport and this facilitates synaptic autophagy, ultimately leading to neurodegeneration, including age-dependent dopaminergic neuron synapse loss. Our work indicates that a fine balance of autophagy at synapses is crucial for neuronal and synapse health and we reveal a Rab39-mediated trafficking mechanism in the neuronal cell body that regulates the sorting of Atg9 vesicles, controlling synaptic autophagy. This cell body-centric process has significant implications for proteostasis at synapses and provides novel insights into the pathophysiology of PD.

## Results

### A forward genetic screen identifies modifiers of autophagy-induced neurodegeneration

To identify regulators of synaptic autophagy, we conducted a forward genetic screen in Drosophila with EndoA^S75D^ mutant flies marked by increased synaptic autophagy and light-induced neurodegeneration (LIND) (Soukup et al. 2016; Bademosi et al. 2023). When these mutant flies are kept in constant light for 3 days a decrease in the depolarization amplitude of the electroretinogram (ERG) response can be observed, indicating LIND (Fig. 1A-B). The ERG is an easy electrophysiological assay that measures the voltage differences in the fly compound eye in response to a 1 s light flash. Degeneration of the neurons in the circuit responding to the light causes a smaller ERG depolarization (Hotta and Benzer 1969). We used this ERG phenotype to conduct a large-scale, unbiased genetic screen to unravel components regulating synaptic autophagy. We generated thousands of EMS-induced lethal mutations on the second chromosome of *Drosophila* (Fig. 1C, labeled P0). This collection was derived from approximately 7000 male flies, each carrying an isogenized second chromosome with *cn bw* mutations (that cause the fly eye to be white – facilitating the assessment of LIND (Escobedo et al. 2022; Tearle 1991). In the subsequent F2 generation, ERGs were recorded from 4403 flies harboring (heterozygous) second chromosomes with a transgene expressing the *endoA^S75D^* mutation and random EMS mutations (Fig. 1C, labeled F2). The presence of 25 EMS-mutated second chromosomes, hereafter called “modifier lines”, rescued the *endoA^S75D^*-induced ERG phenotype (Fig. 1C, F3, Fig. 1D-E).

**Figure 1.**
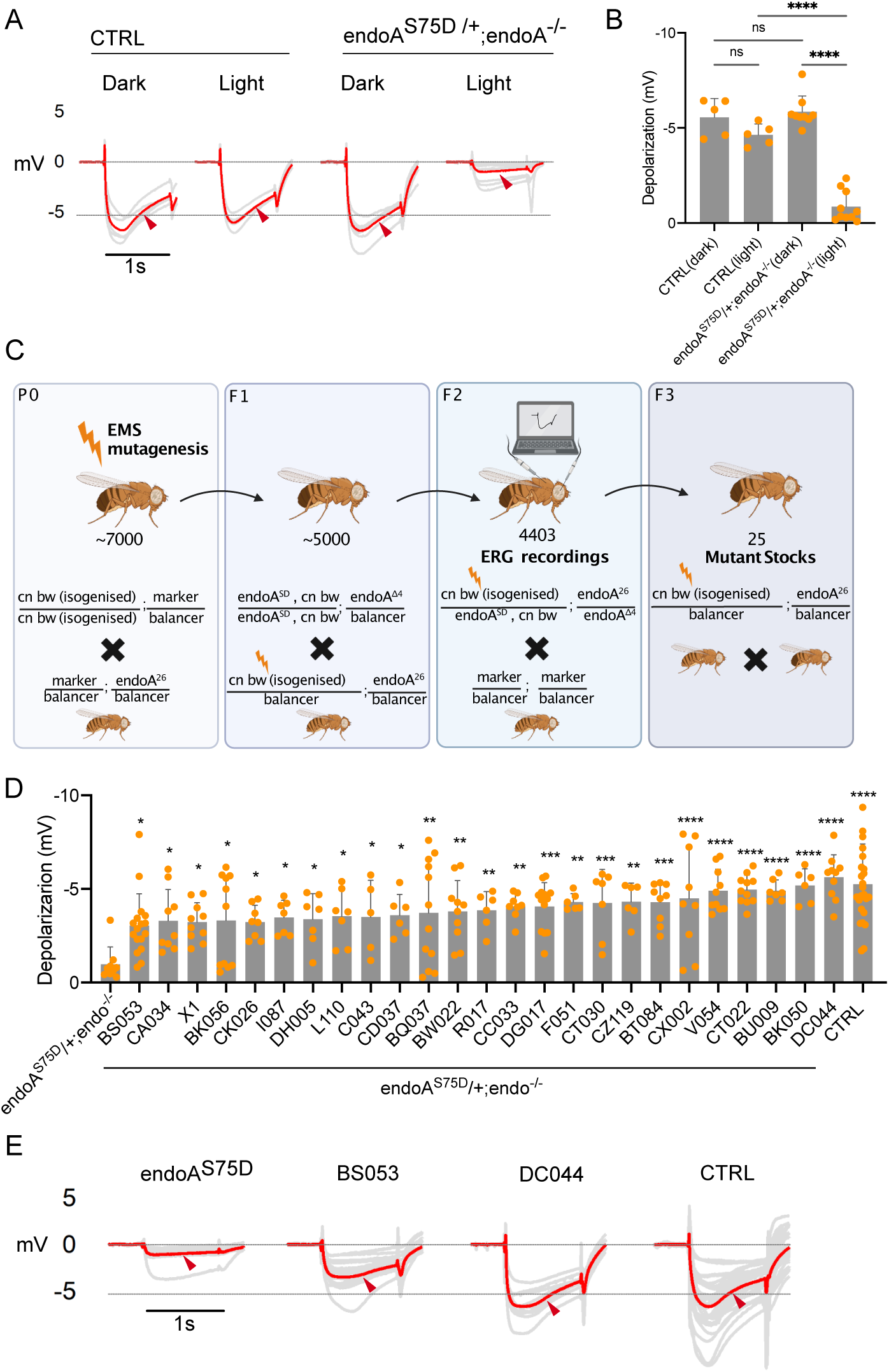
A forward genetic screen identified modifiers of EndoA-mediated neuronal dysfunction. (A) ERG traces of cn bw flies (control, CTRL) and endoA mutant flies expressing endoA^S75D^ in the endoA^-/-^ background (referred as endoA mutant) at endogenous levels (traces of individual flies in gray, average in red). Prior to ERG recordings, animals were exposed to either 3 days of constant light or kept in darkness. Red arrowhead: maximum of depolarization amplitude. (B) Quantification of ERG depolarization amplitudes of flies shown in A. Statistical method: one-way ANOVA with Tukey’s multiple comparison test; n=5-10; error bars: mean ± SD. (C) Schematic overview of the forward genetic screen to identify modifier genes of the EndoA^SD^ light-induced phenotype on the 2nd chromosome using EMS mutagenesis (see methods). (D) Quantifications of the ERG depolarization amplitudes of the 25 mutant lines rescuing the endoA^SD^ phenotype. Statistical method: one-way ANOVA with post hoc Dunnett’s test; n=5-20; error bars: mean ± SD. (E) Example ERG traces of flies carrying modifier mutations (BS053, DC044) that rescued the endoA^SD^-mediated depolarization defect. Mean ERGs were recorded from 5-10 flies kept for 3 days under constant light. Annotated genotypes: cn bw (control, CTRL), endoA^S75D/+^; endoA^-/-^ (endoA^S75D^), BS053/ endoA^S75D^; endoA^-/-^ (BS053), DC044/ endoA^S75D^;endoA^-/-^ (DC044). Red arrowhead: maximum of depolarization amplitude.

### Modifiers of EndoA-induced neurodegeneration also rescue several PD mutants

EndoA interacts with several PD genes (Cao et al. 2014; Watanabe et al. 2018; Soukup et al. 2016) and like EndoA, the PD gene *Rab39* is thought to play a role in autophagy (Corbier and Sellier 2017; Niu et al. 2020) (Fig. 2A). We thus assessed if our modifier lines also affected the phenotypes associated with these other PD genes. When knock-out (KO) mutants of these genes (Kaempf et al. 2024) were exposed to constant light, ERG recordings of *Lrrk^KO-WS^*, *Synj^KO-WS^*, and *Rab39^KO-WS^* (hereafter denoted as “KO”) mutant flies showed a consistent decrease in the amplitude of the depolarization while for *Park^KO-WS^*mutants it did not (Fig. 2B). Interestingly, we found that 12 of the 25 *endoA^S75D^*modifiers also rescued the *Rab39^KO^* ERG phenotype, 6 modifiers rescued the *Synj^KO^* and 5 rescued the *Lrrk^KO^*phenotype (Fig. 2C-D). Hence, several *endoA^S75D^*-modifiers are shared across *Lrrk, Synj* and *Rab39*.

**Figure 2.**
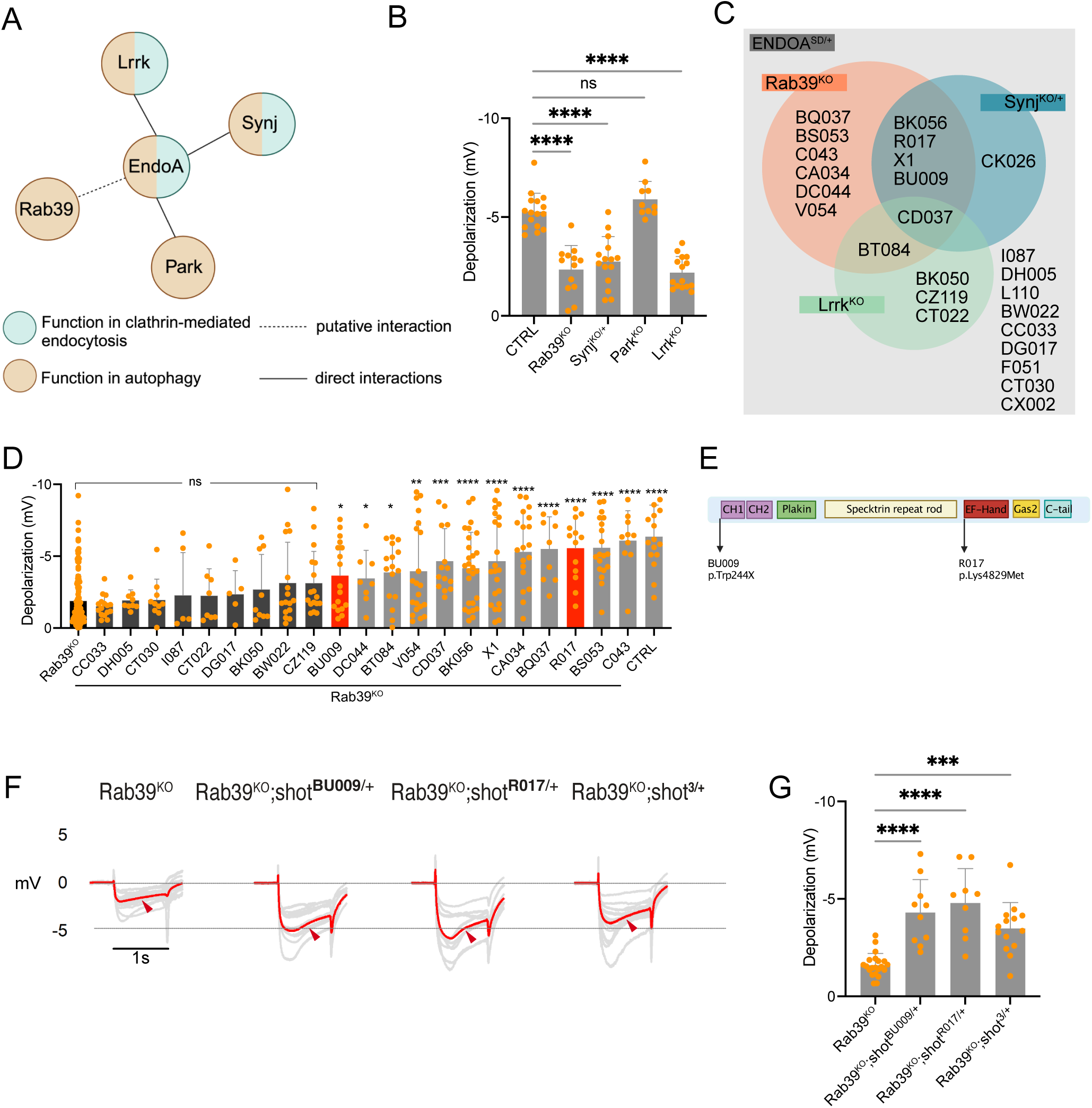
Modifiers of EndoA-induced neurodegeneration are shared among PD mutants. (A) EndoA is at the center of a PD-related functional protein network that links EndoA to three proteins encoded by “PD genes,” Lrrk, Synj, and Park. Proteins functioning in Clathrin-mediated endocytosis are marked in blue, and proteins with a function in (synaptic) autophagy in brown. Direct protein interactions are visualized with a solid line, and putative protein interactions are visualized with a dotted line. (B) Quantification of ERG depolarization amplitude of flies reared under 7 days light of the following genotypes: cn bw (control, CTRL), Rab39^KO^;cn bw, Synj^KO^,cn bw/+, cn bw; Park^KO^, cn bw; Lrrk^KO^. One-way ANOVA followed by Dunnett’s multiple comparisons tests; n=12-16; error bars: mean ± SD. (C) Venn diagram representing the summary ERG data of 25 EMS modifier lines in EndoA^SD^,cn bw;endoA^-/-^, Rab39^KO^,cn bw, Synj^KO^,cn bw/+, cn bw and cn bw, Lrrk^KO^ backgrounds. List of EMS modifier lines rescuing the corresponding mutant. (D) Quantification of the depolarization amplitudes of the screened EMS modifier lines in Rab39^KO^ background. BU009 and R017 lines both harbor a mutation in the shortstop(shot) gene and are indicated in red. One-way ANOVA with post hoc Dunnett’s test; n=5-30; error bars: mean ± SD. (E) Schematic representation of the shot protein showing the location of an amino acid substitution (Lys4829Met) in R017 and a premature stop codon (Trp244X) in BU009 induced by EMS mutagenesis. (F) ERG traces of 7 days light-reared flies with the indicated genotypes. Traces of individual flies in gray, averages in red. Arrowheads show maximum depolarization amplitude. (G) Quantification of the ERG depolarization amplitude of traces shown in (F) Statistical method: one-way ANOVA with Dunnett’s multiple comparisons tests; n=9-20; error bars: mean ± SD.

Given that *endoA^S75D^* and *Rab39^KO^* mutants showed the largest overlap in modifiers, we identified the lesions on the 12 second chromosomes that could rescue both mutants using whole genome sequencing. After filtering the sequence variants (Fig. S1A-B), we retained the mutations that were predicted to have a moderate to high impact on protein function (i.e. mutations affecting splice sites, start and stop codons, and non-synonymous variations) and that occurred more than once independently in the different modifier lines but not in the original isogenized control (Fig. S1A-B, Table S1). This approach yielded 23 genes. GO term analysis using FlyBase biological process annotations indicated a predominant representation of “cytoskeleton organization” (Table S1) (Jenkins, Larkin, and Thurmond 2022). This included a notable hit in *shot* that we recovered independently twice in our screen: *shot^BU009^* and *shot^R017^*. Moreover, a variation at the mammalian *shot* ortholog, *MACF1*, is associated with susceptibility to PD (Wang et al., 2017). Sequencing identified a nonsense mutation at position 244 in *shot^BU009^*, resulting in a premature stop (p.Trp244*), and a missense mutation at position 4829 in *shot^R017^* (p.Lys4829Met) (Fig. 2E). We independently confirmed the causal role for loss of *shot* in rescuing the *Rab39^KO^* ERG phenotype by using the previously generated *shot^3^*null allele (Kolodziej, Yeh Jan, and Nung Jan 1995) that was also effective at rescuing the *Rab39^KO^* ERG defect (Fig. 2F-G). Hence, removing one allele of *shot* rescues *endo^S75D^* and *Rab39^KO^*-induced neurodegeneration.

### *Rab39^KO^* increases synaptic autophagy

To investigate if Rab39, like EndoA, plays a role in synaptic autophagy, we performed live confocal imaging of Atg8^mCherry^ (endogenous expression) at *Rab39^KO^*and control *Drosophila* third-instar larval NMJs under basal conditions. Atg8 is a cytosolic protein that localizes to the autophagosome membrane upon autophagy induction and marks the formation and maturation of autophagosomes (Nakatogawa, Ichimura, and Ohsumi 2007). We observed a significant increase in Atg8-labeled puncta in synaptic boutons of *Rab39^KO^*larvae compared to controls (Fig. 3A and E), and this defect is reduced by re-expression of wild-type Rab39 (*nSyb*-Gal4) (Fig. 3B,D and G). Conversely, we did not observe an increase in Atg8-labeled puncta in neuronal cell bodies, indicating the effect is synapse-specific (Fig. S2A-C, F). The results suggest Rab39 is a negative regulator of synaptic autophagy.

**Figure 3.**
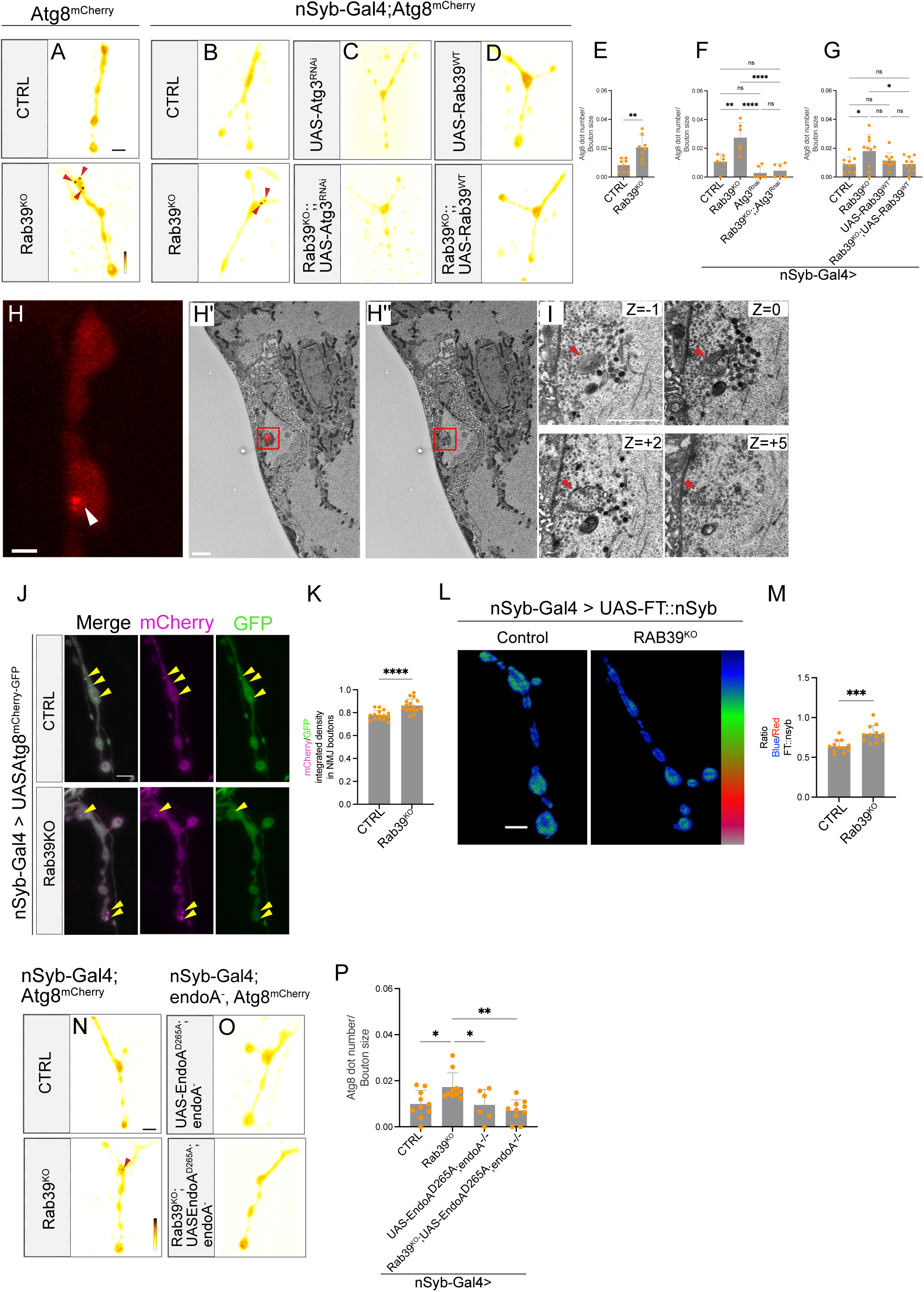
Rab39^KO^ increases synaptic autophagy in *Drosophila* NMJs. (A-D) Live imaging of genomically expressed Atg8^mCherry^ at NMJ boutons of control (W^1118^w+) and Rab39^KO^ (A) and of control and Rab39^KO^ (B) animals expressing UAS-Atg3^RNAi^ (C), UAS-Rab39^WT^ (D) using the nSyb-Gal4 driver. Fluorescence intensities are shown using (423-2667 gray value) indicated in (A) Arrowheads indicate Atg8^mCherry^ puncta. Scale bar: 5 µm. (E-G) Quantification of the number of Atg8^mCherry^ puncta (arrowheads) per NMJ area of the experiment shown in (A-D). Statistical method: Student’s t-test; n= 8 (E); two-way ANOVA with Tukey’s multiple comparison test; n= 6-10 (4 NMJs per animal) (F-G); error bars: mean ± SD. (H-I) CLEM of boutons of Rab39^KO^ animals expressing Atg8^mCherry^. (H) Single confocal plane of an example NMJ displaying an Atg8^mCherry^ puncta (arrowhead). Scale bar: 2 µm. (H’) Overlay of thresholded mCherry positive Atg8 puncta of the bouton in (H) with the TEM image. Scale bar: 2 µm. (H’’) TEM image of the same region as in (H’). (I) Single TEM slices showing the putative autophagosomal structure visible in multiple consecutive slices (red arrowhead). Z = -1: 0, Z = 0: 70 nm, Z = 1: 140 nm, Z = 2: 210 nm, Z = 5: 420 nm. Scale bar: 1 µm. (J) Live imaging of control and Rab39^KO^ larvae expressing UAS-Atg8^mCherry-GFP^ under the control of nSyb-Gal4. mCherry signal in *magenta* and the GFP signal in *green*. Scale bar: 5 µm. Yellow arrowheads mark Atg8 positive puncta. (K) Quantification of the ratio of mCherry signal over GFP integrated density at the NMJ boutons of the experiment shown in (J). Statistical method: Student’s t-test; n=15; error bars: mean ± SD. (L-M) Live imaging of control and Rab39^KO^ animals expressing FT nSyb construct (FT::nSyb) using nSyb-Gal4 in NMJ boutons. Scale bar: 5 µm (L). (M) Quantification of the distribution of the blue (young) over red (old) fluorescence intensities at synaptic boutons analyzed in (L) shown using the indicated lookup table in (L) Statistical method: Student’s t-test; n=12; error bars: mean ± SD. (N-P) Live imaging of genomically expressed Atg8^mCherry^ in NMJ boutons of control (CTRL) and Rab39^KO^ (N) and of endoA^-/-^ animals expressing UAS-EndoA^D265A^ and Rab39^KO^;;endoA^-/-^ animals expressing UAS-EndoA^D265A^ using the nSyb-Gal4 driver. Fluorescence intensities are shown using (904-4095 gray value) indicated in (N). Arrowheads indicate Atg8^mCherry^ puncta. Scale bar: 5 µm. (P) Quantification of the number of Atg8^mCherry^ puncta (arrowheads) per NMJ area of the experiment shown in (N-O). Statistical method: One-way ANOVA with Dunnett’s multiple comparisons tests; n= 6-10 (4 NMJs per animal); error bars: mean ± SD.

We verified that the puncta we observe were indeed autophagosomes by (1) genetics and by (2) imaging: (1) The Atg8^mCherry^ labeled puncta disappeared upon downregulation of the essential autophagy protein Atg3 using Atg3 RNAi in synaptic boutons of control and *Rab39^KO^* larvae (Fig. 3C and F, Fig. S2D-F). (2) We employed correlative light and electron microscopy (CLEM) to visualize the ultrastructural morphology of the mCherry puncta at *Rab39^KO^* synaptic boutons. Alignment of the mCherry fluorescent image with the TEM micrograph showed overlap with structures reminiscent of degradative organelles such as autophagosomes and autolysosomes (Fig. 3H-I).

Finally, we asked if the observed increase in Atg8-positive puncta in *Rab39^KO^* synapses resulted from increased autophagosome biogenesis or a blockage in autophagic flux, and conducted two experiments. We first expressed the dual-tagged autophagic flux reporter Atg8^mCherry-GFP^ in NMJs (Chang and Neufeld 2009). In this assay, Atg8-positive autophagosomes contain both magenta and green fluorescent proteins, resulting in white signal. Following fusion with the acidic lysosome, the GFP signal is quenched by the low pH, leaving only magenta signal (Shinoda, Shannon, and Nagai 2018) (Fig. 3J). At *Rab39^KO^*synapses, we observed a higher magenta over green ratio compared to controls (Fig. 3J-K). This result indicates that *Rab39^KO^* does not block autophagosome acidification and that the mutant either causes increased formation or decreased autophagosome degradation. To distinguish between these possibilities we expressed FT::nSyb, a fluorescent timer (FT) fused to neuronal Synaptobrevin (nSyb) that localizes to synaptic vesicles (SV) (Baumert et al. 1989). The FT converts from blue (young protein) to red (older protein) fluorescence over time. When expressed in *Rab39^KO^* we detected an increased blue over red ratio, indicating increased FT::nSyb turnover (Fig. 3L-M). Together, these data suggest there is increased synaptic autophagic flux in *Rab39^KO^*causing excessive synaptic protein turnover.

Given this result we also assessed the absolute levels of key synaptic proteins such as BRP (Bruchpilot), CSP (Cysteine String Protein), and Syntaxin1A, but found no significant differences between control and *Rab39^KO^*animals (Fig. S3A-C). These proteins play critical roles in synaptic function and neurotransmitter release (Wagh et al. 2006; Umbach, Mastrogiacomo, and Gundersen 1995; Bennett, Calakos, and Scheller 1992), and their stable levels suggest that the excessive autophagy in *Rab39^KO^*does not affect the abundance of all synaptic proteins, including some of the core synaptic machinery, but further work is needed to determine the specific substrates of EndoA-dependent synaptic autophagy and under which conditions these substrates turnover.

### The pathogenic mutant *hRab39B^T168K^* increases synaptic autophagy

Pathogenic mutations in *Rab39B* gene cause PD (Wilson et al. 2014). We therefore determined if the pathogenic human *Rab39B^T168K^* mutant affects synaptic autophagy *in vivo.* We used a knock-in mutant where the first exon of the wildtype fly *Rab39* gene was replaced by either the wildtype human cDNA (*hRab39B*) or a pathogenic T168K mutant variant (*hRab39B^T168K^*) (Pech et al. 2024) and then assessed Atg8^mCherry^ fluorescence at presynaptic terminals under basal conditions. While *hRab39B^WT^*wild type synapses had a similar (low) amount of Atg8-labeled puncta at presynaptic terminals as control animals, this number was significantly higher at *hRab39B^T168K^* mutant synapses (Fig. S4A-E). This result indicates that the pathogenic mutation disregulates autophagy at synapses (as does *LRRK2^G2019S^* (Soukup et al. 2016). Additionally, the data confirm that *hRab39B^T168K^*acts as a loss of Rab39B function (Gao, Martínez-Cerdeño, et al. 2020) and that human Rab39B can compensate for the loss of fly *Rab39*.

### EndoA is epistatic to Rab39

Our data suggest that EndoA and Rab39 have an antagonistic relation regarding synaptic autophagy. We therefore assessed their epistatic relationship. We used the *endoA^D265A^* mutant that blocks synaptic autophagy (Bademosi et al. 2023) and combined it with *Rab39^KO^* that increases synaptic autophagy (*Rab39^KO^; UAS-endoA^D265A^/nSyb-Gal4; endoA^/Δ4^/endoA^26^*). Interestingly, this resulted in low synaptic autophagy levels similar to *endoA^DA^* mutants (Fig. 3N-P) suggesting that EndoA acts downstream of Rab39.

### Rab39 GTPase activity regulates synaptic autophagy

Rab39 is a Rab GTPase alternating between active GTP-bound and inactive GDP-bound states (Müller and Goody 2017) . To determine whether Rab39’s GTPase activity is required to regulate synaptic autophagy, we expressed either wild-type (Rab39^WT^), a constitutively active (Rab39^Q69L^), or a dominant-negative mutant (Rab39^S23N^) in *Rab39^KO^* mutants (Zhang et al. 2007). Rab39^WT^ and the constitutive active Rab39^Q69L^ rescued the elevated synaptic autophagy in *Rab39^KO^* to levels comparable to controls (Fig.4A-D,F). In contrast, overexpressing the inactive Rab39^S23N^ did not rescue the elevated synaptic autophagy levels in *Rab39^KO^* (Fig.4E,F). Hence, Rab39-GTPase activity is required to inhibit synaptic autophagy.

**Figure 4.**
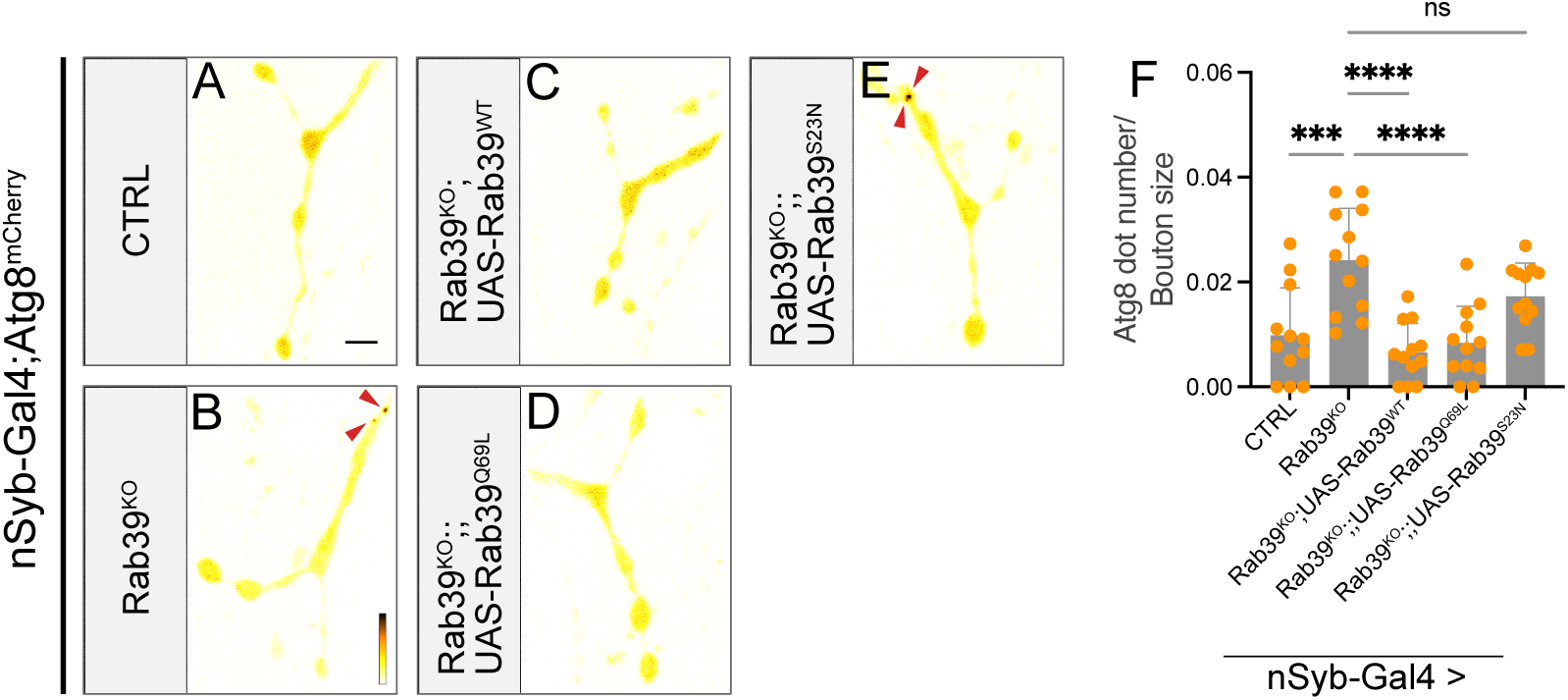
Rab39 GTPase activity regulates synaptic autophagy. (A-D) Live imaging of genomically expressed Atg8^mCherry^ in NMJ boutons of control (W^1118^w+, CTRL) and Rab39^KO^ animals expressing UAS-Rab39^WT^ (C), UAS-Rab39^Q69L^ (D), UAS-Rab39^S23N^ (E) using the nSyb-Gal4 driver. Fluorescence intensities are shown using (571-3514 gray value) indicated in (B) Arrowheads indicate Atg8^mCherry^ puncta. Scale bar 5 µm. (F) Quantification of the number of Atg8^mCherry^ puncta (arrowheads) per NMJ area of the experiment in (A-E). Statistical method: One-way ANOVA with Dunnett’s multiple comparisons tests; n= 12 (4 NMJs per animal); error bars: mean ± SD.

### Rab39 regulates presynaptic autophagy by facilitating the trafficking of Atg9 vesicles

The relationship between EndoA and Rab39 warranted further investigation because the proteins are thought to localize to very different neuronal compartments: EndoA strongly localizes to presynaptic terminals and not to neuronal cell bodies (Verstreken et al. 2002), while Rab39 is thought to be a soma-associated protein (Chan et al. 2011; Gao, Wilson, Stephenson, et al. 2020). We verified this in a line expressing YFP-tagged Rab39 at endogenous levels (Dunst et al. 2015) and indeed found Rab39 to be exclusively located to the neuronal cell body (Fig. S5A-B), without significant labeling at synaptic boutons of NMJs (Fig. S5C). How does a soma-restricted protein control autophagy at synapses?

Our genetic suppressor screen (Fig. 1) had identified regulators of “cytoskeleton organization”, and one of our strongest common suppressors genes was *shot*. Shot is a microtubule-binding protein (Applewhite et al. 2010), stabilizing the axonal cytoskeleton, organizing the axon initial segment (AIS) (Bottenberg et al. 2009; Lee and Kolodziej 2002) and initiating vesicle trafficking in conjunction with the Unc104/Kif1a motor protein (Voelzmann et al. 2016). We reasoned that such a protein could be involved in soma-to-synapse communication to regulate synaptic autophagy. We therefore crossed *shot^3^, shot^BU009^*or *shot^R017^* to *Rab39^KO^*. Heterozygous loss of *shot* not only rescued the *Rab39^KO^*-ERG phenotype, but also restored the levels of presynaptic autophagy (Fig. 5A-G). This was specific to the loss of *shot* function as re-introducing a functional copy of *shot^+^* in *Rab39^KO^*; *shot^3^/+* animals again showed increased synaptic autophagy levels comparable to *Rab39^KO^*(Fig. 5F and 5G). We next examined the interactor of Shot, Unc-104/Kif1a. Heterozygous loss of *unc-104* (*unc-104^P350^*/+) also rescued the increased synaptic autophagy level in *Rab39^KO^* (Fig. 5H-I). Hence, our data suggest that Rab39 regulates synaptic autophagy levels via axonal transport mechanisms.

**Figure 5.**
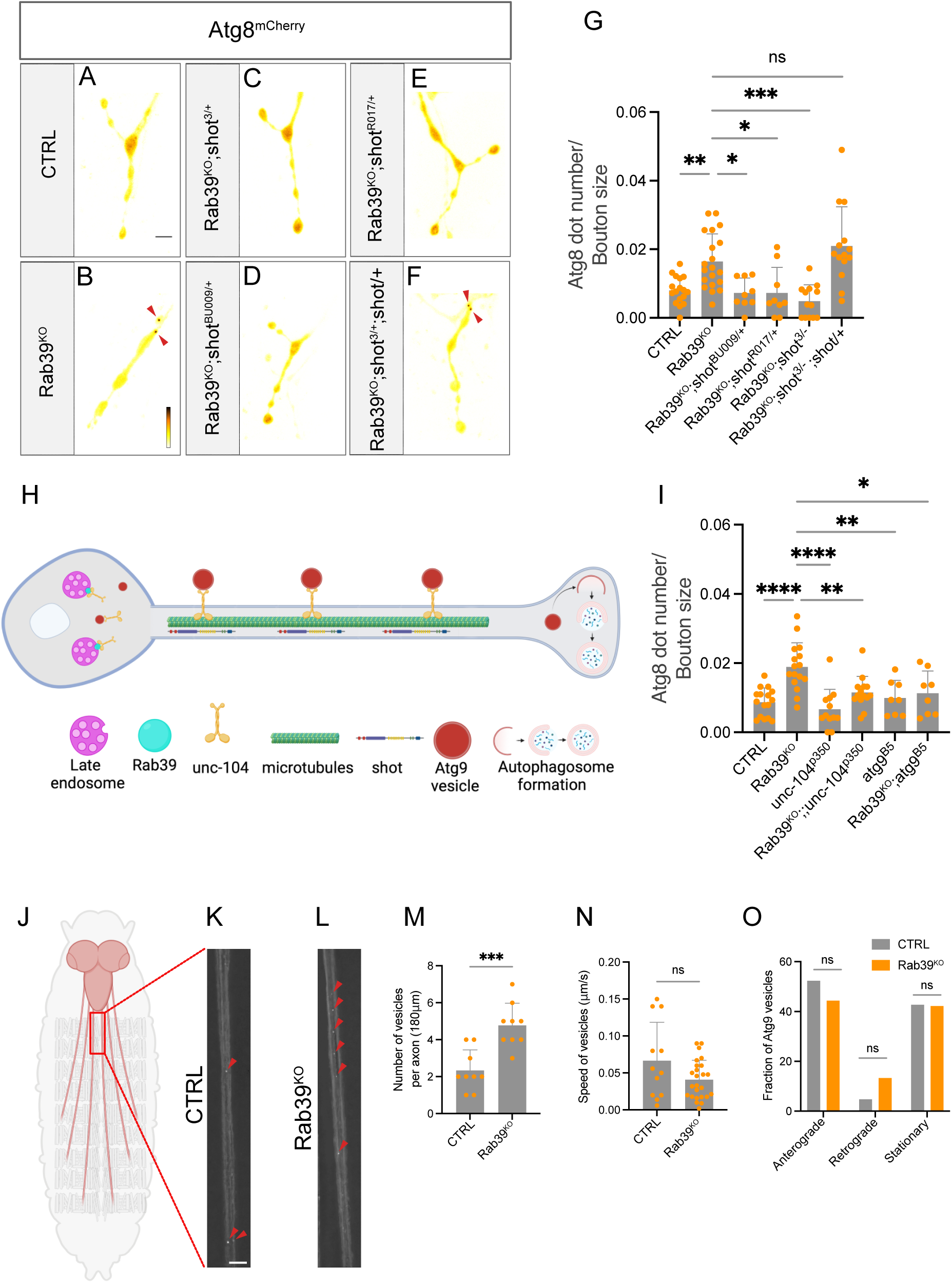
Rab39 regulates presynaptic autophagy by the transport of Atg9 vesicles. (A-F) Live imaging of Atg8^mCherry^ in NMJ boutons of control (W^1118^w+, CTRL) (A), Rab39^KO^ (B), Rab39^KO^;shot^3/+^ (C), Rab39^KO^; shot^BU009/+^ (D), Rab39^KO^; shot^R017/+^ (E), Rab39^KO^;shot^3/+^;shot^+^ (F). Fluorescence intensities are shown using (618-3769 gray value) indicated in (B). Arrowheads indicate Atg8^mCherry^ puncta. Scale bar: 5 µm. (G) Quantification of the number of Atg8^mCherry^ puncta (arrowheads) per NMJ area of (A-F). Statistical method: one-way ANOVA with post hoc Dunnett test; n=9-20 (4 NMJs per animal); error bars: mean ± SD. (H) Schematic model of the proposed function of Rab39 in regulating synaptic autophagy. (I) Quantification of the number of Atg8^mCherry^ puncta (arrowheads) per NMJ area in control and Rab39^KO^ animals carrying null mutants of unc-104 (unc-104^p350^) and atg9 (atg9^B5^). Statistical method: one-way ANOVA with post hoc Dunnett’s test; n=12-16 (4 NMJs per animal); error bars: mean ± SD. (J) A scheme of larval ventral nerve cord and axons in Drosophila 3^rd^ instar larvae indicates the regions imaged for Atg9 vesicle movement analysis. (K-L) Live imaging of control (W^1118^w+, CTRL) (K) and Rab39^KO^ (L) animals expressing genomic Atg9^mcherry^ in atg9^null^ (atg9^B5^) background. Scale bar: 10 µm. Red arrowheads mark Atg9^mCherry^ positive puncta. (M) Quantification of the number of Atg9^mcherry^ puncta in the recorded frame of axons (imaged in K-L) (180 μm) for 180 seconds. Statistical method: Student’s t-test; n= 9; error bars: mean ± SD. (N) Quantification of the speed of moving Atg9^mcherry^ puncta (imaged in K-L). Statistical method: Student’s t-test; n= 12-25; error bars: mean ± SD. (O) Directionality of Atg9 vesicles in control (W^1118^w+, CTRL) and Rab39^KO^ animals (imaged in K-L). Statistical method: Fisher’s exact test.

Atg9 is a transmembrane protein associated with vesicles and it is key for autophagy initiation (Mari et al. 2010; Rao et al. 2016). Such vesicles need to pass the AIS and be transported to synapses to support synaptic autophagy (Stavoe et al. 2016). We first asked if the increased levels of synaptic autophagy in *Rab39^KO^* mutants depended on Atg9 (Fig. 5H-I). While heterozygous loss of Atg9 (*Atg9^B5/+^*) in wildtype larvae does not affect synaptic autophagy, removing a copy of *Atg9* in *Rab39^KO^* (*Rab39^KO^; Atg9^B5/+^*) decreased the elevated synaptic autophagy levels seen in *Rab39^KO^* synaptic boutons (Fig. 5H-I). We then imaged Atg9^mCherry^ transport along motor axons (Fig. 5J). *Rab39^KO^* animals harbored significantly more Atg9^mCherry^-labeled vesicles in their motor axons (Fig. 5K-M). Most Atg9^mCherry^ vesicles travelled anterogradely in both controls and in *Rab39^KO^* larvae and their speed was not affected (Fig. 5N-O). These data are consistent with a model where soma-restricted Rab39 limits the availability of Atg9 vesicles being transported to synapses.

### Intracellular localization of Rab39 changes from late endosome to lysosome upon starvation

It has been shown that Rab39 facilitates trafficking of cargo to lysosomes (Caviglia et al. 2016) . Moreover, Unc-104/Kif1a has been identified as a Rab39 binding partner (Gillingham et al. 2014). Therefore, an appealing hypothesis is that Rab39-mediated lysosomal trafficking regulates Unc-104/Kif1a-dependent Atg9 loading and trafficking. We confirmed by structured illumination-based super-resolution microscopy (SIM) that Rab39 localizes more closely to the endosomal marker Rab7 in neuronal cell bodies. When we acutely induce autophagy by 3-4h starvation (Scott, Schuldiner, and Neufeld 2004) the distance between Rab7 and Rab39 significantly increased (Fig. 6A-D). Conversely, the distance between Rab39 and lysosomes (marked by Lamp1 (Eskelinen, Tanaka, and Saftig 2003)) decreased at cell bodies (Fig. 6E-H). These data confirm Rab39 trafficking between endosomes and lysosomes and they provide evidence this is triggered by starvation. Taken together the data are consistent with a model where autophagy triggers lysosomal delivery of Rab39, possibly leaving Unc104/Kif1a free to be loaded onto Atg9 vesicles for transport.

**Figure 6.**
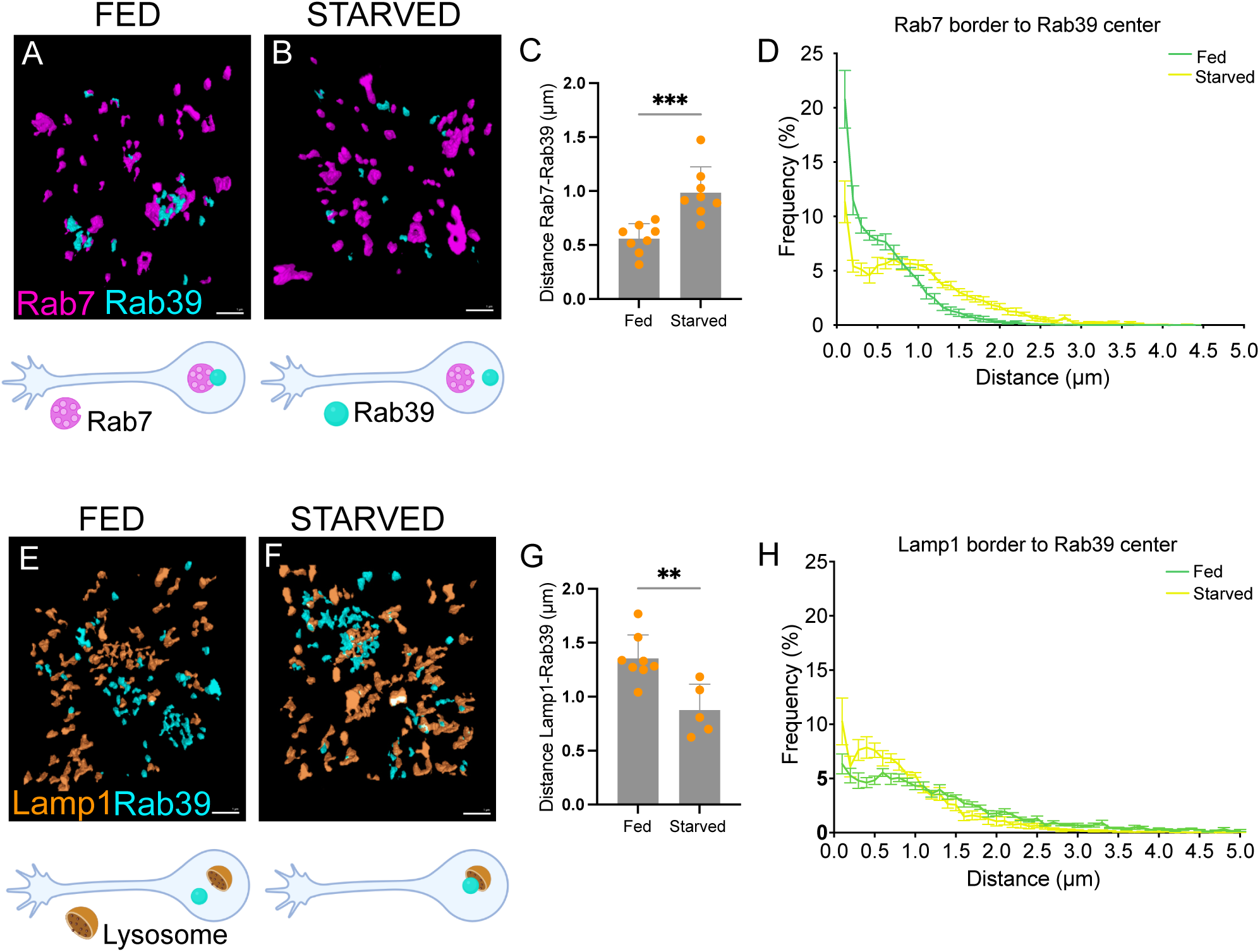
Intercellular localization of Rab39 changes from late endosome to lysosome upon starvation. (A-B) Representative 3D reconstructed images of Rab7 (magenta) and Rab39 (blue) proteins in neuronal cell bodies of the fixed *Drosophila* larval ventral nerve cord in fed (A) and starved conditions (B). Scale bar: 1 µm. (C) Quantification of the distance of the edge of Rab7 objects to the center of the nearest Rab39 object of (A-B). Statistical method: Student’s t-test; n=8; error bars: mean ± SD. (D) The frequency graph of the data quantified from (A-B) of the nearest neighbor distance percentages between Rab7 and Rab39 objects in fed (green) and starved (yellow) conditions. (E-F) Representative 3D reconstructed images of Lamp1 (orange) and Rab39 (blue) proteins in neuronal cell bodies of the fixed *Drosophila* larval ventral nerve cord in fed (E) and starved conditions (F). Scale bar: 1 µm. (G) Quantification of the distance of the edge of Lamp1 objects to the center of the nearest Rab39 objects of (E-F). Statistical method: Student’s t-test; n=5-8; error bars: mean ± SD. (H) The frequency graph of the nearest neighbor distance percentages between Lamp1 and Rab39 objects quantified from (E-F) in fed (green) and starved (yellow) conditions.

### *shot* rescues dopaminergic neuron synapse loss in *Rab39^KO^* mutants

Dopaminergic neuron synapse and cell loss are pathological hallmarks of PD that drive locomotor defects (Antony et al. 2013; Aggarwal, Reichert, and VijayRaghavan 2019). Similarly, *Rab39^KO^* mutant flies suffer from a progressive decline in locomotor behavior and synaptic innervation of protocerebral anterior medial (PAM) dopaminergic neurons (DAN) onto their mushroom bodies (Kaempf et al. 2024; Pech et al. 2024) (Fig 7A). To investigate whether the loss of *shot* function rescues this defect in aging *Rab39^KO^* mutants, DAN and their synapses were labeled using anti-tyrosine hydroxylase (TH) and quantified afferents at the mushroom bodies (labeled by anti-DLG). In young 5-day-old *Rab39^KO^* flies and controls, the mushroom body was well-innervated by TH-positive PAM DAN (Fig. 7B-C’ and Fig. 7J); similarly, *shot^3/+^*heterozygous mutants or *Rab39^KO^; shot^3/+^* showed abundant innervation, similar to controls (Fig. 7D-E’-D). In contrast, 45-day-old *Rab39^KO^*flies show significantly less labeling compared to controls (Fig. 7F-G’ and Fig. 7K). However, this defect was rescued by introducing one copy of *shot^3^* (Fig. 7G-I’ and Fig. 7K). Hence, the loss of *shot* function not only rescues synaptic autophagy in *Rab39* mutants but also the progressive defect of dopaminergic synaptic innervation in the fly brain.

**Figure 7.**
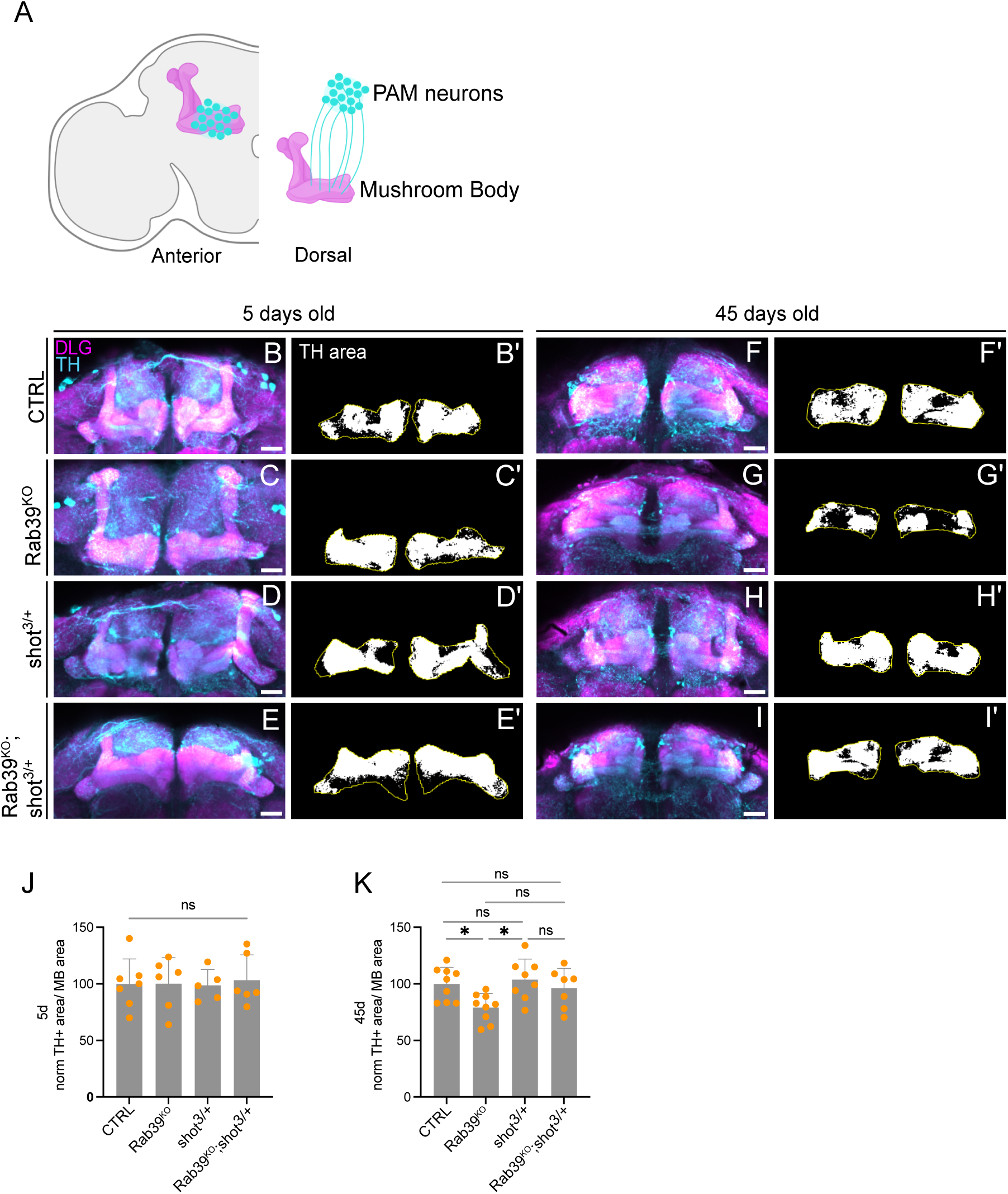
Reducing shot levels partially rescues loss of synapses in TH neurons. (A) Schematic overview of dopaminergic PAM neurons and the mushroom body in anterior (left) and dorsal (right) view of the *Drosophila* adult brain. (B-I’) Maximum intensity projections of confocal images of the mushroom body in the anterior fly brain of 5 days old (B-E’) and 45 days old flies (F-I’), labeled with the dopaminergic marker anti-Tyrosine Hydroxylase (TH) (blue) and the synaptic marker anti-DLG (magenta). (B’-I’) TH-immunoreactive area (white) innervating the mushroom body lobes (yellow ROI) of 5 days old (B’-E’) and 45 days old flies (F’-I’). The anti-DLG area was used to define MB area (yellow outline), anti-TH fluorescence area was quantified as TH+ area. Scale bar: 20 µm (J) Quantification of TH-immunoreactive area (white) within the mushroom body lobes of 5 days old flies in (B-E’). Statistical method: one-way ANOVA with Tukey’s multiple comparison test; n=5-7; error bars: mean ± SD. (K) Quantification of TH-immunoreactive area (white) within the mushroom body lobes of 45 days old flies in (F-I’). Statistical method: one-way ANOVA with Tukey’s multiple comparison test; n=7-9; error bars: mean ± SD.

## Discussion

We demonstrate that Rab39 plays a pivotal role in organizing vesicular trafficking within the neuronal soma towards lysosomes (Seto et al. 2013; Caviglia et al. 2016) *in vivo*, and we propose a model where Rab39 in the cell body regulates the loading of Atg9-positive vesicles onto trafficking motors for transport to synapses. Our data show a significant reduction in Atg9-labeled vesicles within axons of Rab39 mutants, consistent with the hypothesis that Rab39 negatively regulates synaptic autophagy. This suggests a regulatory mechanism active in the soma that governs autophagy at distant synapses.

While synaptic autophagy depends on the activity of several synapse-specific proteins (e.g., BSN, EndoA, Synj1 (Okerlund et al. 2017; Vanhauwaert et al. 2017; Soukup et al. 2016; Nikoletopoulou and Tavernarakis 2018), our findings indicate that central control in the soma remains crucial. We propose that under normal conditions, Rab39 in its GTP-bound state suppresses synaptic autophagy by inhibiting the loading of Atg9-positive vesicles onto trafficking motors. In contrast, when Rab39 transitions to its GDP-bound state, or under pathological conditions, this suppression is relieved. Rab39 interacts with Unc104/Kif1A within a complex, potentially regulated by their shared binding partner, Prd1(Gillingham et al. 2014) . This complex may confer specificity to the loading process of Atg9-positive vesicles.

Although Unc104/Kif1A is a general motor responsible for trafficking diverse cargos to synapses (Hall and Hedgecock 1991; Kern et al. 2013; Pack-Chung et al. 2007), we show that not all synaptic trafficking is impaired in Rab39 mutants. For instance, we observed normal levels of other synapse-enriched proteins (BRP and CSP) and synaptic vesicles, suggesting that Rab39/Prd1/Kif1A may exhibit cargo specificity, with Atg9 vesicles as a particular target. One potential explanation is that active lysosomal trafficking of Rab39GTP-Kif1A complexes limits motor protein availability. When Rab39 is in a GDP-bound state, this constraint may be lifted, allowing motor engagement with Atg9-positive vesicles. This process could be regulated by Prd1, conferring specificity, but future studies are needed to elucidate if Prd1 functions as a linker and regulator in this process.

Our work highlights the critical role of effective axonal transport and microtubule stability in regulating synaptic autophagy. This is supported by our finding that loss-of-function mutations in *shot* and *unc104* rescue the elevated levels of synaptic autophagy observed in *rab39* mutants, and that in our screen we also found other genes with similar functions. Shot and its mammalian orthologs are known to promote microtubule-mediated axonal growth and stability, in part through interactions with EB1, which facilitates axonal growth via microtubule stabilization (Alves-Silva et al. 2012). EB1 also interacts with ankyrin G in the axon initial segment (AIS), a specialized region regulating cargo entry into the axon for further transport (Leterrier et al. 2011). Mammalian Shot orthologs have been shown to bind EB1(Poliakova et al. 2014), suggesting a potential interaction of Shot with the AIS. Interestingly, while we show the trafficking speed and direction of Atg9-positive vesicles are unaffected in *rab39* mutants, our findings suggest that loss of *shot* or *unc104* counteracts excessive Atg9 vesicle trafficking through distinct mechanisms: by removing the motor protein Unc104 or by blocking axonal entry at the AIS via Shot. These genetic manipulations also rescue Rab39-induced neurodegeneration, underscoring the importance of Atg9 vesicle transport and autophagy regulation for neuronal survival.

Our findings connect to observations of deregulated synaptic autophagy in both human and fly models of PD (Decet and Verstreken 2021; Nachman and Verstreken 2022). In the fly central nervous system, both increased and decreased synaptic autophagy lead to neurodegeneration, reflecting the fine balance required for neuronal health. Furthermore pathogenic PD mutations exhibit divergent effects on synaptic autophagy: mutations such as *LRRK2^G2019S^* and, as shown here, *Rab39B^T168K^* increase synaptic autophagy, whereas previous studies have reported that pathogenic mutations in *synj1* and *endoA1* reduce it (Soukup et al. 2016; Vanhauwaert et al. 2017; Bademosi et al. 2023). While familial mutations in these genes are relatively rare, increased levels of alpha-synuclein-a hallmark of both familial and idiopathic Parkinson’s disease (Goedert, Jakes, and Spillantini 2017), are frequently observed and also *postmortem* brain tissue from *Rab39B* mutant patients show extensive Lewy bodies (Wilson et al. 2014; Gao, Martínez-Cerdeño, et al. 2020). Importantly, elevated alpha-synuclein levels correlate with decreased EndoA levels (Westphal and Chandra 2013), and this could in turn again affect synaptic autophagy. These observations imply that the regulation of synaptic autophagy may be a broad phenomenon in idiopathic PD.

The effects of deregulated synaptic autophagy remain unclear. But given that too much and too little of the process are detrimental, one plausible hypothesis is that sensitive signaling systems are disrupted due to impaired autophagy-dependent proteostasis. Future research should focus on identifying the specific targets of this process that contribute to degeneration and determining how to modulate synaptic autophagy to restore and maintain its delicate balance. Such insights could pave the way to understand how PD and related neurodegenerative diseases affect synaptic health and potentially reveal therapeutic inroads.

## Acknowledgements

We thank the Cell and Tissue Imaging Cluster at KU Leuven, supported by Hercules AKUL/11/37 and FWO G.0929.15 to Pieter Vanden Berghe. We also thank Dr. Juhász Gábor for generously providing us with fly stocks. We are grateful to the members of the Verstreken lab for valuable discussions. A.K. (11E2223N), D.V. (151095), and E.M.P. (11O3225N) were supported by PhD fellowships from FWO Vlaanderen, and E.N. (1282123N) was supported by a postdoctoral fellowship from FWO Vlaanderen. Research support was provided in part by the ERC, the Chan Zuckerberg Initiative, a Methusalem grant from the Flemish government, and FWO Vlaanderen. P.V. is an alumnus of the FENS-Kavli Network of Excellence. Cartoons/models/illustrations were created in https://BioRender.com

## Author contributions

Conceptualization, A.K., E.N., V.U., P.V.;

Methodology, A.K., D.V., E.N., V.U., P.V.;

Investigation, A.K., D.V., S.K., J.S., E.M.P., C.C.A., A.E.A., N.C., P.H.V., S.P.;

Formal analysis, A.K., D.V., S.K., A.E.A, S.P.;

Writing, A.K., E.N., V.U, P.V.;

Funding acquisition, A.K., D.V., E.M.P, E.N, P.V.;

Supervision, E.N., V.U., P.V.;

all co-authors read and edited the manuscript.

## Inclusion and diversity

We support inclusive, diverse, and equitable conduct of research.

---

**Figure S1. Filtering process to identify candidate genes in heterozygous mutants**

(A) Flowchart of filters applied to identify candidate mutations in heterozygous EMS* mutants [*cn bw* (*)]. All identified SNVs (ALL) were first filtered against SNVs identified in the isogenized second chromosome (ΔIso). Subsequently, only SNVs that affect the coding sequence or splice sites were retained (functional). Next, the remaining SNVs were filtered against SNVs identified repeatedly in multiple sequenced genomes (ΔEndoA Screen). (B) Impact of filters, introduced in (A), on the total number of SNVs identified on the second chromosome in a 25-Mb interval.

**Figure S2: Autophagy is not affected in the soma of neurons at the ventral nerve cord of Rab39^KO^ mutants**

(A)Representative confocal image of the ventral nerve cord from *Drosophila* larvae expressing Atg8^mCherry^. Scale bar: 100 µm. Fluorescent intensities are shown using (22-2162 gray value). (B-E) Zoomed-in region showing neuronal somas of control (CTRL) (B), Rab39^KO^ (C), UAS-Atg3^Rnai^ (D), and Rab39^KO^;;Atg3^Rnai^ (E) flies driven by nSyb-Gal4. Scale bar: 10 µm. Fluorescent intensities are shown using (89-609 gray value). Neuronal cell bodies are thresholded and outlined in red, mCherry puncta are marked for particle counting. (F) Quantification of Atg8^mCherry^ puncta per cell area of the experiment in (B-E). Statistical method: one-way ANOVA with Tukey’s multiple comparisons test; n = 12; error bars represent mean ± SD.

**Figure S3. Synaptic protein levels in control and Rab39^KO^ animals**

(A-C) Representative maximum projection confocal images of NMJ boutons of control (W^1118^w+, CTRL) and Rab39^KO^ animals stained with anti-HRP (magenta) to outline the presynaptic NMJ and with anti-BRP labeling active zones (green) (A), with anti-CSP labeling synaptic vesicle-associated protein (green) (B), with anti-Syntaxin-1A labeling a membrane-associated protein at presynaptic sites (green) (C). Scale bar: 5 µm. (A’-C’) Quantification of BRP (A’), CSP (B’) and Syntaxin-1A (C’) average pixel intensity per NMJ area (HRP-positive area). Statistical method: Student’s t-test; n=5; error bars: mean ± SD.

**Figure S4. Pathogenic Rab39B^T168K^ mutant mimics Rab39^KO^ and increases synaptic autophagy**

(A-D) Live imaging of genomically expressed Atg8^mCherry^ in NMJ boutons of Control (W^1118^w+, CTRL) (A), Rab39^KO^ (B), hRab39B (C) and hRab39B^T168K^ (D). Red arrowheads indicate Atg8^mCherry^ puncta. Scale bar 5: µm. (E) Quantification of the number of Atg8^mCherry^ puncta (arrowheads) per NMJ area od (A-D). Statistical method: two-way ANOVA with Tukey’s multiple comparison test; n= 8; Error bars: mean ± SD.

**Figure S5. Localization of Rab39^EYFP^ at the cell bodies of the ventral nerve cord and at NMJs**

(A)Airyscan confocal image of the larval ventral nerve cord at 20X magnification, stained with anti-GFP (green) to visualize endogenously tagged Rab39^EYFP^ and anti-DLG (magenta) to label postsynaptic regions. Scale bar: 50 µm. (B) Higher magnification (63X) of the ventral nerve cord region shown in (A) Scale bar 20 µm. (C) High-magnification (63X) image of NMJ bouton stained with anti-GFP and anti-DLG, showing the localization of Rab39^EYFP^ within NMJ boutons. Scale bar:10 µm.

## METHODS

### Resource availability

#### Lead Contact

Further information and requests for resources and reagents should be directed to the lead contact, Patrik Verstreken.

#### Materials Availability

Data, code, Drosophila models, and reagents are available upon request.

### *Drosophila* lines

Fruit flies were maintained in the incubator at 25°C on a standard diet containing cornmeal and molasses with a 12:12 light-dark cycle. CRISPR/Cas9-mediated gene editing was employed to generate a collection of *Drosophila* PD models, as described in (Pech et al. 2024; Kaempf et al. 2024). Briefly, each knockout line had an attP-flanked w+ reporter cassette inserted to replace the first common exon across isoforms of the targeted gene. In this study, we used synj, lrrk, rab39, and park knockout flies from this collection (Kaempf et al. 2024). The control line, w^1118^ was further modified by inserting a transgenic mini-white marker on the X-chromosome, resulting in a red-eyed control (w^1118^ w+). The following genotypes were used in this study:

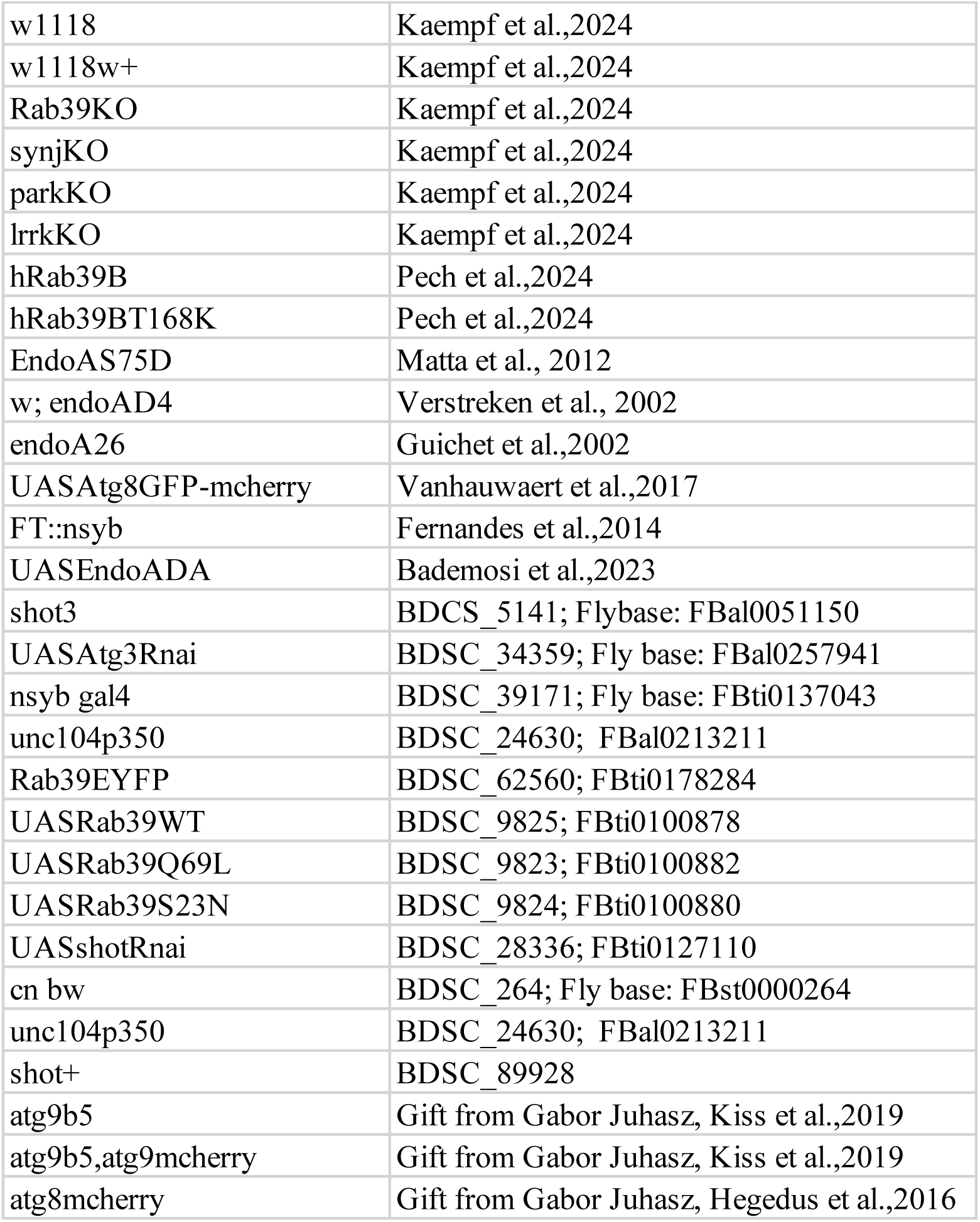

### Ethyl methanesulfonate (EMS) mutagenesis

Mutagenesis was performed on isogenized cn bw^iso^ males that were fed 15 nM EMS in a sucrose solution. After recovery from mutagenesis, these males were mated with endoA null virgin females. In the F1 generation, cn bw mut*;endoA null (mut* indicated the EMS-induced mutation) virgins were collected and crossed with endoA^SD^;endoA null males to conduct ERG phenotype screening. 4403 lines carrying lethal mutations were balanced and kept, the remaining stocks were discarded.

### Light-induced neurodegeneration and ERGs

All ERG experiments were conducted with male flies in a *cn bw* background to deplete eye pigment; these flies have white eyes. For ERG recordings, flies were collected 1 after eclosion and kept in the incubator at 25°C until day 3. Three-day-old male flies were transferred to an incubator where they were exposed to constant illumination (1300 LUX, 24h/day) at 25°C. EndoA^SD^ mutant and EMS-generated mutants used for ERG screening were kept under constant light for 3 days, while PD knock-out mutants, including Rab39^KO^, Lrrk^KO^, Synj^KO^ and Park^KO^ were maintained under continuous light exposure for 7 days. For the ERG measurements, flies were immobilized on glass microscope slides with double-sided tape. Filamented glass micropipettes were used to generate electrodes using the Laser-pipette-puller P-2000 (Sutter Instrument). Electrodes were filled with a 3 M NaCl solution. The recording electrode was placed lightly onto the ommatidia. The reference electrode was inserted into the thorax. The flies were exposed to darkness for 3 s, followed by 1 s of LED light illumination. This was repeated 5 times for each fly. Light-evoked signals were amplified by a DC amplifier and the amplified signal was processed by a data acquisition device (Clampex), connected to a PC running Clampfit software. ERG traces were analyzed in IGOR Pro 6.37 using a custom-made macro.

### Next-Generation Sequencing and candidate gene selection

Next-generation sequencing of flies was performed at the VIB Nucleomics Core as described in the Illumina NebNEXT DNA Ultra protocol (Gautreau 2015). The GATK4 (v4.0.9.0) Germline short variant discovery (SNPs + Indels) workflow (Poplin et al. 2017) was adapted to the Drosophila genome and applied to the Illumina paired read data. Paired reads were aligned to the *Drosophila* reference genome (Dm_BDGP6) (Hoskins et al. 2015). Known variants were obtained from NCBI (Sherry et al. 2001) and FlyVar (F. Wang et al. 2015). The final VCF data was annotated, filtered with SnpEff (v4.3) (Cingolani et al. 2012) and internally developed logic detailed below to obtain various candidate lists detailed in the supplementary figure S1. FlyBase gene ontology terms were used for gene ontology analysis (Jenkins, Larkin, and Thurmond 2022).

### Confocal Live Imaging and quantification

Third instar larvae expressing Atg8mCherry, FT-nSyb or Atg8^mCherry-GFP^ were dissected in fresh HL3 (100 mM NaCl, 5 mM KCl, 10 mM NaHCO_3_, 5 mM Hepes, 30 mM sucrose, 5 mM trehalose, 10 mM MgCl_2_, pH 7.2). Live imaging of dissected larvae was performed using a Nikon NiE A1R confocal microscope equipped with a 60X (NA 1.0) water-dipping objective. Prior to imaging, the dissected larvae were rinsed several times in HL3 solution. Data acquisition was conducted through NIS-Elements (Nikon) software, employing the Resonant Scanning mode, with a zoom of 3 and 16-fold line averaging. Images were collected with a pinhole setting of one Airy unit and a resolution of 1024 x 1024 pixels. Z-stacks were taken to capture fluorescence (puncta) throughout the entire NMJ. Puncta within synaptic boutons were quantified using ImageJ.15, with Atg8^mCherry^ puncta manually counted and a threshold mask applied to define bouton areas. Trafficking autophagosomes (puncta in motor neuron axons outside the NMJ) were excluded from the analysis.

To image the FTs, blue and red channels were acquired (excitation: blue, 405 nm and red, 561 nm). For ratio calculations, the intensities of the boutons were quantified for both blue and red channels, and the ratios were calculated. The ratio images were prepared using the FIJI image processing package: both channels were first converted to a 32-bit image and thresholded, retaining only the bouton labeling; then, the “red” image was divided by the “blue” using the arithmetic module, and a look-up table was applied to the resulting image. As a result, the images only include the bouton areas (the thresholded area), and the rest of the image was intentionally left black.

To perform the Atg8^mCherry-GFP^ tandem experiment to measure autophagic flux, green and red channels were used to image (excitation: green, 488 nm and red, 561 nm). For ratio calculations, the intensities of the boutons were quantified for both green and red channels, and the ratios were calculated.

### Correlative Light Electron Microscopy (CLEM)

To characterize the Atg8-mCherry-positive structures with their ultrastructure, CLEM was performed (Bademosi et al. 2023; Vanhauwaert et al. 2017; Soukup et al. 2016; Decet et al. 2024) . First, third instar *Rab39^KO^* mutant larvae endogenously expressing Atg8^mCherry^ were dissected in cold Ca^2+^ free HL3 (100 mM NaCl, 5 mM KCl, 10 mM NaHCO_3_, 5 mM Hepes, 30 mM sucrose, 5 mM trehalose, 10 mM MgCl_2_, pH 7.2) and subsequently fixed for 2 h at 4°C (0.5 % glutaraldehyde, 2 % paraformaldehyde in 0.1 M phosphate buffer (PB), pH 7.4). After washing in 0.1 M PB, fillets were incubated with DAPI (Sigma). Next, using a Zeiss LSM 780 equipped with a Mai Tai HP DeepSee laser (Spectra-Physics) at 880 nm with 40 % maximal power output, near-infrared branding (NIRB) was used to apply visible branding marks. Before and after branding, Z stacks of the region of interest (ROI) were acquired with a 25 X water immersion lens (NA 0.8). Immediately after NIRB, fillets were post-fixed (4 % paraformaldehyde, 2.5 % glutaraldehyde in 0.1 PB) overnight at 4°C and washed with 0.1 M PB and ddH_2_O until the dehydration steps. Subsequently, fillets were first osmicated for 1 h (1 % OsO_4_ and 1.5 % potassium ferrocyanide) and then incubated in a 0.2 % tannic acid for 30 min followed by a second osmication step (1 % OsO_4_ for 30 min) and next incubated for 20 min in 1 % thiocarbohydrazide. Again, fillets were osmicated for a third time (1 % OsO_4_ for 30 min) and incubated overnight in 0.5 % uranyl acetate. Thereafter, fillets were stained with lead aspartate (Walton’s lead aspartate: 20 mM lead nitrate in 30 mM sodium aspartate, pH 5.5) for 30 min at 60°C. After a final washing step, and a dehydration series (with solutions of increasing ethanol concentration (30 %, 50 %, 70 %, 90 % and twice with 100 %)), fillets were incubated twice for 10 min with propylene oxide. Finally, fillets were infiltrated with resin agar 100 (Laborimpex), flat embedded in resin agar 100 and placed at 60°C for 48 h.

The flat resin-embedded samples were cropped into 1 mm^2^ pieces with ROI in the middle and sectioned until the first branding marks were reached and muscle morphology was recognized by correlating with the light microscopy data. Next, ultrathin sections (70 nm) were cut on an ultramicrotome (EM UC7, Leica), collected on 1×2 mm slot, copper grids (Ted Pella, inc) and imaged using a JEM-1400 transmission electron microscope (Jeol) at 80 keV. NIRB branding marks around the NMJ and DAPI signal were used to correlate the confocal images with the TEM micrographs of the NMJ boutons. Overlay images were generated using ImageJ and GIMP.

### Immunohistochemistry

For immunohistochemistry of larvae samples, dissections were performed in HL3 (100 mM NaCl, 5 mM KCl, 10 mM NaHCO_3_, 5 mM Hepes, 30 mM sucrose, 5 mM trehalose, 10 mM MgCl_2_, pH 7.2) by opening the third instar larvae dorsally along the midline and removing the entrails on the sylgard plate.

For NMJ stainings, the larval fillets were fixated with 3.7 % paraformaldehyde in HL3 for 20 minutes. Fixed larvae were permeabilized with 0.4 % PBX (TritonX-100 in 1X PBS) 4 times for 15 minutes on a shaker, blocked for 1 hour with 10 % normal goat serum in PBX and incubated overnight at 4°C with primary antibodies. After several washes with 0.4 % PBX, larval filets were incubated with secondary antibodies for 90 min at room temperature. Samples were mounted in Vectashield (Vector Laboratories).

For ventral nerve cord stainings, larval fillet was fixated with Bouin’s fixative for 10 min. After fixation, the filets were permeabilized with 0.05% PBX 4 times for 15 minutes on a shaker and blocked for 1 h in 5% normal goat serum in PBX and incubated overnight at 4°C with primary antibodies. After several washes, larval filets were incubated with secondary antibodies overnight at 4°C. Samples were mounted in RapiClear 1.47 (SUNJin Lab) on 1.5H high performance coverslip (Marienfeld). To induce starvation to image Rab39, Rab7 and Lamp1 localization in VNC, third-instar larvae were placed in petri dishes with 20% sucrose and 1% agarose for 3–4 hours before dissection. The following antibodies were used: rabbit anti-HRP [1:2000 (Jackson Immuno Research)], mouse anti-DLG [1:50 (DSHB)], mouse anti-Syx1A(8C3) [1:20 (DSHB)], mouse anti-Brp(nc82) [1:250 (DSHB)], chicken anti-GFP [1:500 (Invitrogen)], rabbit anti-Lamp1[1:500 (Abcam)], mouse anti-Rab7 [1:50 (DSHB)] and rabbit anti-Rab39B [1:500, proteintech], Alexa Fluor-488/Alexa Fluor-555 conjugated secondary antibodies [2:500 (Invitrogen)].

For immunohistochemistry of adult brain samples, brains of 5-and 45-days old animals were dissected in ice-cold PBS and fixed for 30 minutes in a freshly prepared 3.7% paraformaldehyde solution containing 1X PBS and 0.2% Triton X-100 (PBX) at room temperature (RT). After three 15-minute washes in PBX at RT on a shaker, brains were placed in a blocking solution (PBX with 10% normal goat serum (NGS)) for one hour at RT. The samples were then incubated with primary antibodies (rabbit anti-TH from Sigma at 1:200 and mouse anti-DLG from DSHB at 1:100) in blocking solution at 4°C for 1.5-2 days, followed by three 15-minute washes in PBX at RT. Secondary antibodies (goat anti-rabbit Alexa488 and goat anti-mouse Alexa555 at 1:500) in PBX with 10% NGS were applied overnight at 4°C. Afterward, brains were washed three times for 15 minutes in PBT at RT on a shaker and then mounted anterior-side up in RapiClear 1.47 (SUNJin Lab). Entire brains were imaged as Z-stacks using a Nikon A1R confocal microscope with a 20x (NA 0.95) water immersion objective.

### Structured illumination microscopy imaging

The Localization of Rab39^EYFP^ with Rab7 and Lamp1 was imaged by a Nikon Ti2 N-SIM S microscope equipped with a Hamamatsu ORCA-Flash 4.0 (C13440) in combination with an SR HP Plan Apo Lambda S 100x Sil lens (NA 1.35) with silicon immersion oil. The set up was controlled by NIS-Elements 5.30.07 (Build 1569). Stacks of 2 µm or 4 µm were acquired in 3D-SIM mode with a step size of 0.1 µm. Samples were excited sequentially either with 488 nm or 561 nm laser and emission was recorded with a bandpass filter 525/30 and 595/30, respectively. Image analysis was done in batch with NIS-Elements 5.42.03 (Nikon). The GA3 analysis protocol included background subtraction of a constant (noise background), 3D threshold, filter by size and measure distances from the center of Rab39 objects to border of organelle object. Image visualization was done using Arrivis Vision 4D (version 4.12, Arivis AG)

### Airyscan Confocal Imaging

The localization of Rab39^EYFP^ in the larval ventral nerve cord and neuromuscular junctions (NMJs) was imaged using a Zeiss LSM900 microscope equipped with an Airyscan2 module. Imaging was performed with a 63X oil objective lens (NA 1.4) and a 20X air objective lens (NA 0.80). ZEN blue (3.7, Carl Zeiss Microscopy GmbH) software was used for image acquisition and processing.

### Spinning Disc Imaging and quantification

For the tracking of Atg9 vesicles, images were acquired using a Nikon NiE upright microscope equipped with a Yokogawa CSU-X spinning-disk module and a Prime BSI camera (Teledyne Photometrics) in combination with a 60X water dipping objective (numerical aperture 1.0). The system was controlled by NIS-Elements software (5.42.06, Nikon Instruments Europe B.V.). Live imaging was performed in the TRITC channel (excitation wavelength 561 nm, emission wavelength 595 nm) at a resolution of 1192 x 1192 pixels (mono, 12-bit). Time-lapse images were recorded at 10-second intervals over a total duration of 3 minutes. Each time-lapse series included 21 Z-stacks taken with a step size of 1 μm. Post-processing and image analysis were conducted in NIS-Elements (version 5.42.03, Nikon). Images were pre-processed in batch using the GA3 analysis protocol. Vesicles were manually traced within the NIS-Elements software. The “Overlay” setting was enabled with arrows to mark the tracked path, a new ROI was defined by selecting the circular ROI around each vesicle. Each vesicle was identified at the starting point of the time-lapse and tracked across frames by dragging the marker to follow its movement in each subsequent frame. Track binaries tool is used to calculate the speed and the heading direction of the vesicles.

### Confocal imaging and quantification

Adult brain images were imaged as Z-stacks using a Nikon NiE A1R confocal microscope with a 20X (NA 0.95) water immersion objective. Z-stacks were collected with 3 μm step intervals, maintaining identical image settings across genotypes and imaging sessions. Image analysis was performed in Fiji. Dopaminergic neuron innervation of the mushroom body (MB) was quantified by identifying the five Z-stacks containing the synaptic region of the MB lobes using anti-DLG. The MB region of interest (ROI) was defined in a sum projection of these five Z-planes, and within these, anti-TH fluorescence areas were thresholded using a uniform threshold to exclude background signal similar to the control. The thresholded area in the ROI was measured in each z-plane, summed, and normalized to the MB area for each brain individually (TH+ area/MB area). In each experiment, individual TH+ area/MB area values were normalized to the control mean. For aged fly images, the maximum projection of five Z-planes and a thresholded middle Z-plane are shown.

### Statistics

Statistical significance was determined using GraphPad Prism 10. Significance levels used were as follows:

****p < 0.0001, ***p < 0.001, **p < 0.01,*p < 0.05, and ns = p > 0.05.

## References

Aggarwal, Aman, Heinrich Reichert, and K. VijayRaghavan. 2019. “A Locomotor Assay Reveals Deficits in Heterozygous Parkinson’s Disease Model and Proprioceptive Mutants in Adult *Drosophila*.” Proceedings of the National Academy of Sciences 116 (49): 24830–39. 10.1073/pnas.1807456116.

Alves-Silva, Juliana, Natalia Sánchez-Soriano, Robin Beaven, Melanie Klein, Jill Parkin, Thomas H. Millard, Hugo J. Bellen, et al. 2012. “Spectraplakins Promote Microtubule-Mediated Axonal Growth by Functioning as Structural Microtubule-Associated Proteins and EB1-Dependent +TIPs (Tip Interacting Proteins).” The Journal of Neuroscience: The Official Journal of the Society for Neuroscience 32 (27): 9143–58. 10.1523/JNEUROSCI.0416-12.2012.

Antony, Paul M. A., Nico J. Diederich, Rejko Krüger, and Rudi Balling. 2013. “The Hallmarks of Parkinson’s Disease.” The FEBS Journal 280 (23): 5981–93. 10.1111/febs.12335.

Applewhite, Derek A., Kyle D. Grode, Darby Keller, Alireza Dehghani Zadeh, Kevin C. Slep, and Stephen L. Rogers. 2010. “The Spectraplakin Short Stop Is an Actin–Microtubule Cross-Linker That Contributes to Organization of the Microtubule Network.” Edited by Paul Forscher. Molecular Biology of the Cell 21 (10): 1714–24. 10.1091/mbc.e10-01-0011.

Bademosi, Adekunle T., Marianna Decet, Sabine Kuenen, Carles Calatayud, Jef Swerts, Sandra F. Gallego, Nils Schoovaerts, et al. 2023. “EndophilinA-Dependent Coupling between Activity-Induced Calcium Influx and Synaptic Autophagy Is Disrupted by a Parkinson-Risk Mutation.” Neuron 111 (9): 1402–1422.e13. 10.1016/j.neuron.2023.02.001.

Baumert, M., P. R. Maycox, F. Navone, P. De Camilli, and R. Jahn. 1989. “Synaptobrevin: An Integral Membrane Protein of 18,000 Daltons Present in Small Synaptic Vesicles of Rat Brain.” The EMBO Journal 8 (2): 379–84. 10.1002/j.1460-2075.1989.tb03388.x.

Bennett, M. K., N. Calakos, and R. H. Scheller. 1992. “Syntaxin: A Synaptic Protein Implicated in Docking of Synaptic Vesicles at Presynaptic Active Zones.” *Science (New York*, N.Y.) 257 (5067): 255–59. 10.1126/science.1321498.

Birdsall, Veronica, and Clarissa L. Waites. 2019. “Autophagy at the Synapse.” Neuroscience Letters 697 (April):24–28. 10.1016/j.neulet.2018.05.033.

Bottenberg, Wolfgang, Natalia Sanchez-Soriano, Juliana Alves-Silva, Ines Hahn, Michael Mende, and Andreas Prokop. 2009. “Context-Specific Requirements of Functional Domains of the Spectraplakin Short Stop in Vivo.” Mechanisms of Development 126 (7): 489–502. 10.1016/j.mod.2009.04.004.

Cao, Mian, Ira Milosevic, Silvia Giovedi, and Pietro De Camilli. 2014. “Upregulation of Parkin in Endophilin Mutant Mice.” Journal of Neuroscience 34 (49): 16544–49. 10.1523/JNEUROSCI.1710-14.2014.

Caviglia, Sara, Marko Brankatschk, Elisabeth J. Fischer, Suzanne Eaton, and Stefan Luschnig. 2016. “Staccato/Unc-13-4 Controls Secretory Lysosome-Mediated Lumen Fusion during Epithelial Tube Anastomosis.” Nature Cell Biology 18 (7): 727–39. 10.1038/ncb3374.

Chan, Chih-Chiang, Shane Scoggin, Dong Wang, Smita Cherry, Todd Dembo, Ben Greenberg, Eugene Jennifer Jin, et al. 2011. “Systematic Discovery of Rab GTPases with Synaptic Functions in Drosophila.” Current Biology 21 (20): 1704–15. 10.1016/j.cub.2011.08.058.

Chang, Diana, Mike A. Nalls, Ingileif B. Hallgrímsdóttir, Julie Hunkapiller, Marcel van der Brug, Fang Cai, International Parkinson’s Disease Genomics Consortium, et al. 2017. “A Meta-Analysis of Genome-Wide Association Studies Identifies 17 New Parkinson’s Disease Risk Loci.” Nature Genetics 49 (10): 1511–16. 10.1038/ng.3955.

Chang, Yu-Yun, and Thomas P. Neufeld. 2009. “An Atg1/Atg13 Complex with Multiple Roles in TOR-Mediated Autophagy Regulation.” Molecular Biology of the Cell 20 (7): 2004–14. 10.1091/mbc.E08-12-1250.

Cingolani, Pablo, Adrian Platts, Le Lily Wang, Melissa Coon, Tung Nguyen, Luan Wang, Susan J. Land, Xiangyi Lu, and Douglas M. Ruden. 2012. “A Program for Annotating and Predicting the Effects of Single Nucleotide Polymorphisms, SnpEff.” Fly 6 (2): 80–92. 10.4161/fly.19695.

Corbier, Camille, and Chantal Sellier. 2017. “C9ORF72 Is a GDP/GTP Exchange Factor for Rab8 and Rab39 and Regulates Autophagy.” Small GTPases 8 (3): 181–86. 10.1080/21541248.2016.1212688.

Decet, Marianna, Patrick Scott, Sabine Kuenen, Douja Meftah, Jef Swerts, Carles Calatayud, Sandra F. Gallego, et al. 2024. “A Candidate Loss-of-Function Variant in SGIP1 Causes Synaptic Dysfunction and Recessive Parkinsonism.” Cell Reports Medicine 5 (10): 101749. 10.1016/j.xcrm.2024.101749.

Decet, Marianna, and Patrik Verstreken. 2021. “Presynaptic Autophagy and the Connection With Neurotransmission.” Frontiers in Cell and Developmental Biology 9 (December). 10.3389/fcell.2021.790721.

Dou, Dan, Jayne Aiken, and Erika L.F. Holzbaur. 2024. “RAB3 Phosphorylation by Pathogenic LRRK2 Impairs Trafficking of Synaptic Vesicle Precursors.” The Journal of Cell Biology 223 (6): e202307092. 10.1083/jcb.202307092.

Dunst, Sebastian, Tom Kazimiers, Felix von Zadow, Helena Jambor, Andreas Sagner, Beate Brankatschk, Ali Mahmoud, et al. 2015. “Endogenously Tagged Rab Proteins: A Resource to Study Membrane Trafficking in Drosophila.” Developmental Cell 33 (3): 351–65. 10.1016/j.devcel.2015.03.022.

Escobedo, Spencer E., Sarah C. Stanhope, Ziyu Dong, and Vikki M. Weake. 2022. “Aging and Light Stress Result in Overlapping and Unique Gene Expression Changes in Photoreceptors.” Genes 13 (2): 264. 10.3390/genes13020264.

Eskelinen, Eeva-Liisa, Yoshitaka Tanaka, and Paul Saftig. 2003. “At the Acidic Edge: Emerging Functions for Lysosomal Membrane Proteins.” Trends in Cell Biology 13 (3): 137–45. 10.1016/s0962-8924(03)00005-9.

Fernandes, Ana Clara, Valerie Uytterhoeven, Sabine Kuenen, Yu Chun Wang, Jan R. Slabbaert, Jef Swerts, Jaroslaw Kasprowicz, Stein Aerts, and Patrik Verstreken. 2014. “Reduced Synaptic Vesicle Protein Degradation at Lysosomes Curbs TBC1D24/Sky-Induced Neurodegeneration.” Journal of Cell Biology 207 (4): 453–62. 10.1083/jcb.201406026.

Fujimoto, Tetta, Tomoki Kuwahara, Tomoya Eguchi, Maria Sakurai, Tadayuki Komori, and Takeshi Iwatsubo. 2018. “Parkinson’s Disease-Associated Mutant LRRK2 Phosphorylates Rab7L1 and Modifies Trans-Golgi Morphology.” Biochemical and Biophysical Research Communications 495 (2): 1708–15. 10.1016/j.bbrc.2017.12.024.

Gao, Yujing, Verónica Martínez-Cerdeño, Kirk J. Hogan, Catriona A. McLean, and Paul J. Lockhart. 2020. “Clinical and Neuropathological Features Associated With Loss of RAB39B.” Movement Disorders: Official Journal of the Movement Disorder Society 35 (4): 687–93. 10.1002/mds.27951.

Gao, Yujing, Gabrielle R. Wilson, Nicholas Salce, Alexandra Romano, George D. Mellick, Sarah E.M. Stephenson, and Paul J. Lockhart. 2020. “Genetic Analysis of RAB39B in an Early-Onset Parkinson’s Disease Cohort.” Frontiers in Neurology 11 (June). 10.3389/fneur.2020.00523.

Gao, Yujing, Gabrielle R. Wilson, Sarah E.M. Stephenson, Mustapha Oulad-Abdelghani, Nicolas Charlet- Berguerand, Kiymet Bozaoglu, Catriona A. McLean, Paul Q. Thomas, David I. Finkelstein, and Paul J. Lockhart. 2020. “Distribution of Parkinson’s Disease Associated RAB39B in Mouse Brain Tissue.” Molecular Brain 13 (1). 10.1186/s13041-020-00584-7.

Gautreau, Isabel. 2015. “NEBNext® Ultra^TM^ DNA Library Prep Protocol for Illumina® With Size Selection (E7370),” February. https://www.protocols.io/view/NEBNext-Ultra-DNA-Library-Prep-Protocol-for-Illumi-imsty5.

Gillingham, Alison K., Rita Sinka, Isabel L. Torres, Kathryn S. Lilley, and Sean Munro. 2014. “Toward a Comprehensive Map of the Effectors of Rab GTPases.” Developmental Cell 31 (3): 358–358. 10.1016/j.devcel.2014.10.007.

Goedert, Michel, Ross Jakes, and Maria Grazia Spillantini. 2017. “The Synucleinopathies: Twenty Years On.” Journal of Parkinson’s Disease 7 (s1): S51–69. 10.3233/JPD-179005.

Grosso Jasutkar, Hilary, and Ai Yamamoto. 2023. “Autophagy at the Synapse, an Early Site of Dysfunction in Neurodegeneration.” Current Opinion in Physiology 32 (April):100631. 10.1016/j.cophys.2023.100631.

Guichet, Antoine, Tanja Wucherpfennig, Veronica Dudu, Sylvain Etter, Michaela Wilsch-Bräuniger, Andrea Hellwig, Marcos González-Gaitán, Wieland B. Huttner, and Anne A. Schmidt. 2002. “Essential Role of Endophilin A in Synaptic Vesicle Budding at the Drosophila Neuromuscular Junction.” The EMBO Journal 21 (7): 1661–72. 10.1093/emboj/21.7.1661.

Gustavsson, Emil K, Jordan Follett, Joanne Trinh, Sandeep K Barodia, Raquel Real, Zhiyong Liu, Melissa Grant-Peters, et al. 2024. “RAB32 Ser71Arg in Autosomal Dominant Parkinson’s Disease: Linkage, Association, and Functional Analyses.” The Lancet Neurology 23 (6): 603–14. 10.1016/S1474-4422(24)00121-2.

Hall, David H., and Edward M. Hedgecock. 1991. “Kinesin-Related Gene *Unc-104* Is Required for Axonal Transport of Synaptic Vesicles in C. Elegans.” Cell 65 (5): 837–47. 10.1016/0092-8674(91)90391-B.

Hegedűs, Krisztina, Szabolcs Takáts, Attila Boda, András Jipa, Péter Nagy, Kata Varga, Attila L. Kovács, and Gábor Juhász. 2016. “The Ccz1-Mon1-Rab7 Module and Rab5 Control Distinct Steps of Autophagy.” Edited by Suresh Subramani. Molecular Biology of the Cell 27 (20): 3132–42. 10.1091/mbc.e16-03-0205.

Hop, Paul J., Dongbing Lai, Pamela J. Keagle, Desiree M. Baron, Brendan J. Kenna, Maarten Kooyman, Shankaracharya, et al. 2024. “Systematic Rare Variant Analyses Identify RAB32 as a Susceptibility Gene for Familial Parkinson’s Disease.” Nature Genetics 56 (7): 1371–76. 10.1038/s41588-024-01787-7.

Hoskins, Roger A., Joseph W. Carlson, Kenneth H. Wan, Soo Park, Ivonne Mendez, Samuel E. Galle, Benjamin W. Booth, et al. 2015. “The Release 6 Reference Sequence of the Drosophila Melanogaster Genome.” Genome Research 25 (3): 445–58. 10.1101/gr.185579.114.

Hotta, Yoshiki, and Seymour Benzer. 1969. “Abnormal Electroretinograms in Visual Mutants of Drosophila.” Nature 222 (5191): 354–56. 10.1038/222354a0.

Jacobson, Jessie R., Capucine Piat, Allen J. Aksamit, Gaia Patane’, Owen A. Ross, and Rodolfo Savica. 2024. “Novel RAB39B Loss-of-Function Mutation in Patient with Typical Early-Onset Parkinson’s Disease.” Parkinsonism & Related Disorders 123 (June). 10.1016/j.parkreldis.2024.106038.

Jenkins, Victoria K., Aoife Larkin, and Jim Thurmond. 2022. “Using FlyBase: A Database of Drosophila Genes and Genetics.” In Drosophila: Methods and Protocols, edited by Christian Dahmann, 1–34. New York, NY: Springer US. 10.1007/978-1-0716-2541-5_1.

Kaempf, Natalie, Jorge S. Valadas, Pieter Robberechts, Nils Schoovaerts, Roman Praschberger, Antonio Ortega, Ayse Kilic, et al. 2024. “Behavioral Screening Defines Three Molecular Parkinsonism Subgroups in Drosophila.” bioRxiv. 10.1101/2024.08.27.609924.

Kern, Jeannine V, Yao V Zhang, Stella Kramer, Jay E Brenman, and Tobias M Rasse. 2013. “The Kinesin-3, Unc-104 Regulates Dendrite Morphogenesis and Synaptic Development in *Drosophila*.” Genetics 195 (1): 59–72. 10.1534/genetics.113.151639.

Kia, Demis A., David Zhang, Sebastian Guelfi, Claudia Manzoni, Leon Hubbard, Regina H. Reynolds, Juan Botía, et al. 2021. “Identification of Candidate Parkinson Disease Genes by Integrating Genome-Wide Association Study, Expression, and Epigenetic Data Sets.” JAMA Neurology 78 (4): 464–72. 10.1001/jamaneurol.2020.5257.

Kiss, Viktória, András Jipa, Kata Varga, Szabolcs Takáts, Tamás Maruzs, Péter Lőrincz, Zsófia Simon- Vecsei, et al. 2020. “Drosophila Atg9 Regulates the Actin Cytoskeleton via Interactions with Profilin and Ena.” Cell Death and Differentiation 27 (5): 1677–92. 10.1038/s41418-019-0452-0.

Kolodziej, Peter A., Lily Yeh Jan, and Yuh Nung Jan. 1995. “Mutations That Affect the Length, Fasciculation, or Ventral Orientation of Specific Sensory Axons in the Drosophila Embryo.” Neuron 15 (2): 273–86. 10.1016/0896-6273(95)90033-0.

Lee, Seungbok, and Peter A. Kolodziej. 2002. “Short Stop Provides an Essential Link between F-Actin and Microtubules during Axon Extension.” Development 129 (5): 1195–1204. 10.1242/dev.129.5.1195.

Leterrier, Christophe, Hélène Vacher, Marie-Pierre Fache, Stéphanie Angles d’Ortoli, Francis Castets, Amapola Autillo-Touati, and Bénédicte Dargent. 2011. “End-Binding Proteins EB3 and EB1 Link Microtubules to Ankyrin G in the Axon Initial Segment.” Proceedings of the National Academy of Sciences 108 (21): 8826–31. 10.1073/pnas.1018671108.

Liu, Zhiyong, Nicole Bryant, Ravindran Kumaran, Alexandra Beilina, Asa Abeliovich, Mark R. Cookson, and Andrew B. West. 2018. “LRRK2 Phosphorylates Membrane-Bound Rabs and Is Activated by GTP-Bound Rab7L1 to Promote Recruitment to the Trans-Golgi Network.” Human Molecular Genetics 27 (2): 385–95. 10.1093/hmg/ddx410.

Mari, Muriel, Janice Griffith, Ester Rieter, Lakshmi Krishnappa, Daniel J. Klionsky, and Fulvio Reggiori. 2010. “An Atg9-Containing Compartment That Functions in the Early Steps of Autophagosome Biogenesis.” The Journal of Cell Biology 190 (6): 1005–22. 10.1083/jcb.200912089.

Matta, Samer, Kristof Van Kolen, Raquel da Cunha, Geert van den Bogaart, Wim Mandemakers, Katarzyna Miskiewicz, Pieter Jan De Bock, et al. 2012. “LRRK2 Controls an EndoA Phosphorylation Cycle in Synaptic Endocytosis.” Neuron 75 (6): 1008–21. 10.1016/j.neuron.2012.08.022.

Müller, Matthias P., and Roger S. Goody. 2017. “Molecular Control of Rab Activity by GEFs, GAPs and GDI.” Small GTPases 9 (1–2): 5. 10.1080/21541248.2016.1276999.

Nachman, Eliana, and Patrik Verstreken. 2022. “Synaptic Proteostasis in Parkinson’s Disease.” Current Opinion in Neurobiology 72 (February):72–79. 10.1016/j.conb.2021.09.001.

Nakatogawa, Hitoshi, Yoshinobu Ichimura, and Yoshinori Ohsumi. 2007. “Atg8, a Ubiquitin-like Protein Required for Autophagosome Formation, Mediates Membrane Tethering and Hemifusion.” Cell 130 (1): 165–78. 10.1016/j.cell.2007.05.021.

Nalls, Mike A., Cornelis Blauwendraat, Costanza L. Vallerga, Karl Heilbron, Sara Bandres-Ciga, Diana Chang, Manuela Tan, et al. 2019. “Identification of Novel Risk Loci, Causal Insights, and Heritable Risk for Parkinson’s Disease: A Meta-Genome Wide Association Study.” The Lancet. Neurology 18 (12): 1091–1102. 10.1016/S1474-4422(19)30320-5.

Nikoletopoulou, Vassiliki, and Nektarios Tavernarakis. 2018. “Regulation and Roles of Autophagy at Synapses.” Trends in Cell Biology 28 (8): 646–61. 10.1016/j.tcb.2018.03.006.

Niu, Mengxi, Naizhen Zheng, Zijie Wang, Yue Gao, Xianghua Luo, Zhicai Chen, Xing Fu, et al. 2020. “RAB39B Deficiency Impairs Learning and Memory Partially Through Compromising Autophagy.” Frontiers in Cell and Developmental Biology 8 (December). 10.3389/fcell.2020.598622.

Okerlund, Nathan D., Katharina Schneider, Sergio Leal-Ortiz, Carolina Montenegro-Venegas, Sally A. Kim, Loren C. Garner, Clarissa L. Waites, Eckart D. Gundelfinger, Richard J. Reimer, and Craig C. Garner. 2017. “Bassoon Controls Presynaptic Autophagy through Atg5.” Neuron 93 (4): 897–913.e7. 10.1016/j.neuron.2017.01.026.

Pack-Chung, Eunju, Peri T Kurshan, Dion K Dickman, and Thomas L Schwarz. 2007. “A Drosophila Kinesin Required for Synaptic Bouton Formation and Synaptic Vesicle Transport.” NATURE NEUROSCIENCE 10 (8): 10.

Pech, Ulrike, Jasper Janssens, Nils Schoovaerts, Sabine Kuenen, Samira Makhzami, Gert Hulselmans, Suresh Poovathingal, et al. 2024. “Synaptic Deregulation of Cholinergic Projection Neurons Causes Olfactory Dysfunction across 5 Fly Parkinsonism Models.” eLife 13 (July). 10.7554/eLife.98348.1.

Poliakova, Kseniia, Adijat Adebola, Conrad L. Leung, Bertrand Favre, Ronald K. H. Liem, Isabelle Schepens, and Luca Borradori. 2014. “BPAG1a and b Associate with EB1 and EB3 and Modulate Vesicular Transport, Golgi Apparatus Structure, and Cell Migration in C2.7 Myoblasts.” Edited by Yanmin Yang. PLoS ONE 9 (9): e107535. 10.1371/journal.pone.0107535.

Poplin, Ryan, Valentin Ruano-Rubio, Mark A. DePristo, Tim J. Fennell, Mauricio O. Carneiro, Geraldine A. Van Der Auwera, David E. Kling, et al. 2017. “Scaling Accurate Genetic Variant Discovery to Tens of Thousands of Samples,” November. 10.1101/201178.

Purlyte, Elena, Herschel S. Dhekne, Adil R. Sarhan, Rachel Gomez, Pawel Lis, Melanie Wightman, Terina N. Martinez, Francesca Tonelli, Suzanne R. Pfeffer, and Dario R. Alessi. 2018. “Rab29 Activation of the Parkinson’s Disease-Associated LRRK2 Kinase.” The EMBO Journal 37 (1): 1–18. 10.15252/embj.201798099.

Rao, Yijian, Marco G. Perna, Benjamin Hofmann, Viola Beier, and Thomas Wollert. 2016. “The Atg1– Kinase Complex Tethers Atg9-Vesicles to Initiate Autophagy.” Nature Communications 7 (1): 10338. 10.1038/ncomms10338.

Schimmöller, Frauke, Iris Simon, and Suzanne R. Pfeffer. 1998. “Rab GTPases, Directors of Vesicle Docking *.” Journal of Biological Chemistry 273 (35): 22161–64. 10.1074/jbc.273.35.22161.

Schirinzi, Tommaso, Graziella Madeo, Giuseppina Martella, Marta Maltese, Barbara Picconi, Paolo Calabresi, and Antonio Pisani. 2016. “Early Synaptic Dysfunction in Parkinson’s Disease: Insights from Animal Models.” Movement Disorders: Official Journal of the Movement Disorder Society 31 (6): 802–13. 10.1002/mds.26620.

Scott, Ryan C., Oren Schuldiner, and Thomas P. Neufeld. 2004. “Role and Regulation of Starvation-Induced Autophagy in the *Drosophila* Fat Body.” Developmental Cell 7 (2): 167–78. 10.1016/j.devcel.2004.07.009.

Seto, Shintaro, Keiko Sugaya, Kunio Tsujimura, Toshi Nagata, Toshinobu Horii, and Yukio Koide. 2013. “Rab39a Interacts with Phosphatidylinositol 3-Kinase and Negatively Regulates Autophagy Induced by Lipopolysaccharide Stimulation in Macrophages.” PLoS ONE 8 (12). 10.1371/journal.pone.0083324.

Sherry, S. T., M.-H. Ward, M. Kholodov, J. Baker, L. Phan, E. M. Smigielski, and K. Sirotkin. 2001. “dbSNP: The NCBI Database of Genetic Variation.” Nucleic Acids Research 29 (1): 308. 10.1093/nar/29.1.308.

Shinoda, Hajime, Michael Shannon, and Takeharu Nagai. 2018. “Fluorescent Proteins for Investigating Biological Events in Acidic Environments.” International Journal of Molecular Sciences 19 (6): 1548. 10.3390/ijms19061548.

Song, Pingping, Wesley Peng, Veronique Sauve, Rayan Fakih, Zhong Xie, Daniel Ysselstein, Talia Krainc, et al. 2023. “Parkinson’s Disease-Linked Parkin Mutation Disrupts Recycling of Synaptic Vesicles in Human Dopaminergic Neurons.” Neuron 111 (23): 3775–3788.e7. 10.1016/j.neuron.2023.08.018.

Soukup, Sandra Fausia, Sabine Kuenen, Roeland Vanhauwaert, Julia Manetsberger, Sergio Hernández-Díaz, Jef Swerts, Nils Schoovaerts, et al. 2016. “A LRRK2-Dependent EndophilinA Phosphoswitch Is Critical for Macroautophagy at Presynaptic Terminals.” Neuron 92 (4): 829–44. 10.1016/j.neuron.2016.09.037.

Stavoe, Andrea K.H., Sarah E. Hill, David H. Hall, and Daniel A. Colón-Ramos. 2016. “KIF1A/UNC-104 Transports ATG-9 to Regulate Neurodevelopment and Autophagy at Synapses.” Developmental Cell 38 (2): 171–85. 10.1016/j.devcel.2016.06.012.

Steger, Martin, Francesca Tonelli, Genta Ito, Paul Davies, Matthias Trost, Melanie Vetter, Stefanie Wachter, et al. 2016. “Phosphoproteomics Reveals That Parkinson’s Disease Kinase LRRK2 Regulates a Subset of Rab GTPases.” Edited by Ivan Dikic. eLife 5 (January):e12813. 10.7554/eLife.12813.

Tan, Manuela M. X., Michael A. Lawton, Miriam I. Pollard, Emmeline Brown, Raquel Real, Alejandro Martinez Carrasco, Samir Bekadar, et al. 2024. “Genome-Wide Determinants of Mortality and Motor Progression in Parkinson’s Disease.” Npj Parkinson’s Disease 10 (1): 1–15. 10.1038/s41531-024-00729-8.

Tearle, Rick. 1991. “Tissue Specific Effects of Ommochrome Pathway Mutations in Drosophila Melanogaster.” Genetics Research 57 (3): 257–66. 10.1017/S0016672300029402.

Umbach, J. A., A. Mastrogiacomo, and C. B. Gundersen. 1995. “Cysteine String Proteins and Presynaptic Function.” Journal of Physiology, Paris 89 (2): 95–101. 10.1016/0928-4257(96)80556-0.

Vanhauwaert, Roeland, Sabine Kuenen, Roy Masius, Adekunle Bademosi, Julia Manetsberger, Nils Schoovaerts, Laura Bounti, et al. 2017. “The SAC 1 Domain in Synaptojanin Is Required for Autophagosome Maturation at Presynaptic Terminals.” The EMBO Journal 36 (10): 1392–1411. 10.15252/embj.201695773.

Verstreken, Patrik, Ole Kjaerulff, Thomas E. Lloyd, Richard Atkinson, Yi Zhou, Ian A. Meinertzhagen, and Hugo J. Bellen. 2002. “Endophilin Mutations Block Clathrin-Mediated Endocytosis but Not Neurotransmitter Release.” Cell 109 (1): 101–12. 10.1016/S0092-8674(02)00688-8.

Vijayan, Vinoy, and Patrik Verstreken. 2017. “Autophagy in the Presynaptic Compartment in Health and Disease.” Journal of Cell Biology 216 (7): 1895–1906. 10.1083/jcb.201611113.

Voelzmann, Andre, Pilar Okenve-Ramos, Yue Qu, Monika Chojnowska-Monga, Manuela del Caño-Espinel, Andreas Prokop, and Natalia Sanchez-Soriano. 2016. “Tau and Spectraplakins Promote Synapse Formation and Maintenance through Jun Kinase and Neuronal Trafficking.” eLife 5 (August). 10.7554/elife.14694.

Wagh, Dhananjay A., Tobias M. Rasse, Esther Asan, Alois Hofbauer, Isabell Schwenkert, Heike Dürrbeck, Sigrid Buchner, et al. 2006. “Bruchpilot, a Protein with Homology to ELKS/CAST, Is Required for Structural Integrity and Function of Synaptic Active Zones in Drosophila.” Neuron 49 (6): 833–44. 10.1016/j.neuron.2006.02.008.

Wang, Fei, Lichun Jiang, Yong Chen, Nele A. Haelterman, Hugo J. Bellen, and Rui Chen. 2015. “FlyVar: A Database for Genetic Variation in Drosophila Melanogaster.” Database 2015. 10.1093/database/bav079.

Wang, Xin, Nuomin Li, Nian Xiong, Qi You, Jie Li, Jinlong Yu, Hong Qing, et al. 2017. “Genetic Variants of Microtubule Actin Cross-Linking Factor 1 (MACF1) Confer Risk for Parkinson’s Disease.” Molecular Neurobiology 54 (4): 2878–88. 10.1007/s12035-016-9861-y.

Watanabe, Shigeki, Lauren Elizabeth Mamer, Sumana Raychaudhuri, Delgermaa Luvsanjav, Julia Eisen, Thorsten Trimbuch, Berit Söhl-Kielczynski, et al. 2018. “Synaptojanin and Endophilin Mediate Neck Formation during Ultrafast Endocytosis.” Neuron 98 (6): 1184–1197.e6. 10.1016/j.neuron.2018.06.005.

Westphal, Christopher H., and Sreeganga S. Chandra. 2013. “Monomeric Synucleins Generate Membrane Curvature.” The Journal of Biological Chemistry 288 (3): 1829–40. 10.1074/jbc.M112.418871.

Wilson, Gabrielle R., Joe C.H. Sim, Catriona McLean, Maila Giannandrea, Charles A. Galea, Jessica R. Riseley, Sarah E.M. Stephenson, et al. 2014. “Mutations in RAB39B Cause X-Linked Intellectual Disability and Early-Onset Parkinson Disease with α-Synuclein Pathology.” American Journal of Human Genetics 95 (6): 729–35. 10.1016/j.ajhg.2014.10.015.

Yang, Sisi, Daehun Park, Laura Manning, Sarah E. Hill, Mian Cao, Zhao Xuan, Ian Gonzalez, et al. 2022. “Presynaptic Autophagy Is Coupled to the Synaptic Vesicle Cycle via ATG-9.” Neuron 110 (5): 824–840.e10. 10.1016/j.neuron.2021.12.031.

Zhang, Jun, Karen L Schulze, P Robin Hiesinger, Kaye Suyama, Stream Wang, Matthew Fish, Melih Acar, Roger A Hoskins, Hugo J Bellen, and Matthew P Scott. 2007. “Thirty-One Flavors of Drosophila Rab Proteins.” Genetics 176 (2): 1307–22. 10.1534/genetics.106.066761.

