## Supplementary figures for "Soma-centered control of synaptic autophagy by Rab39-regulated anterograde trafficking of Atg9"

**Figure S1. Filtering process to identify candidate genes in heterozygous mutants**

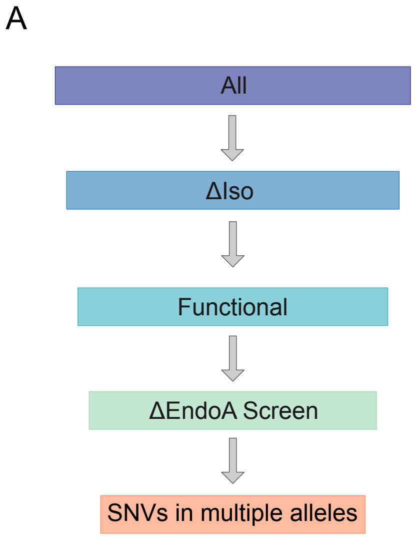


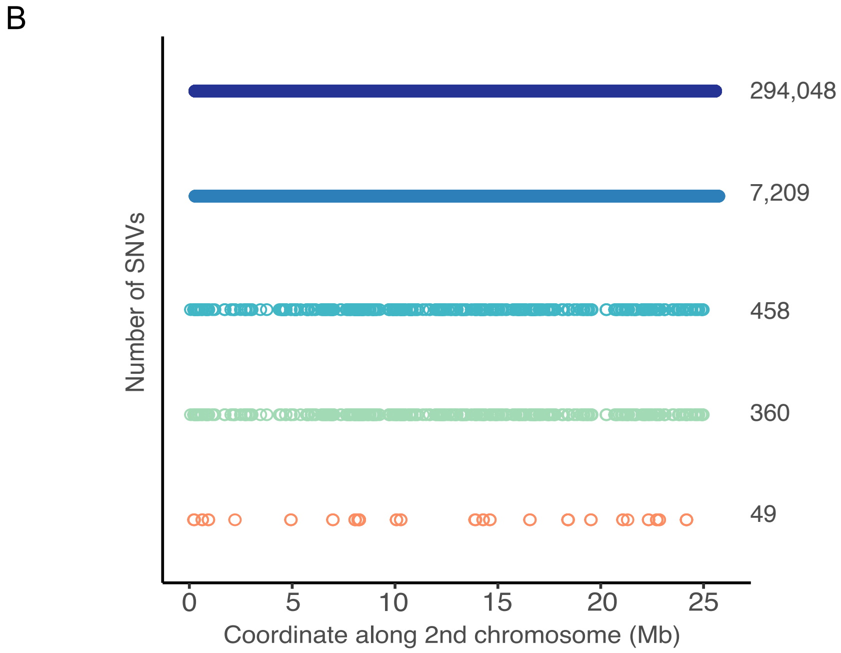

**Figure S2. Autophagy is not affected in the soma of neurons at the ventral nerve cord of Rab39KO mutants**

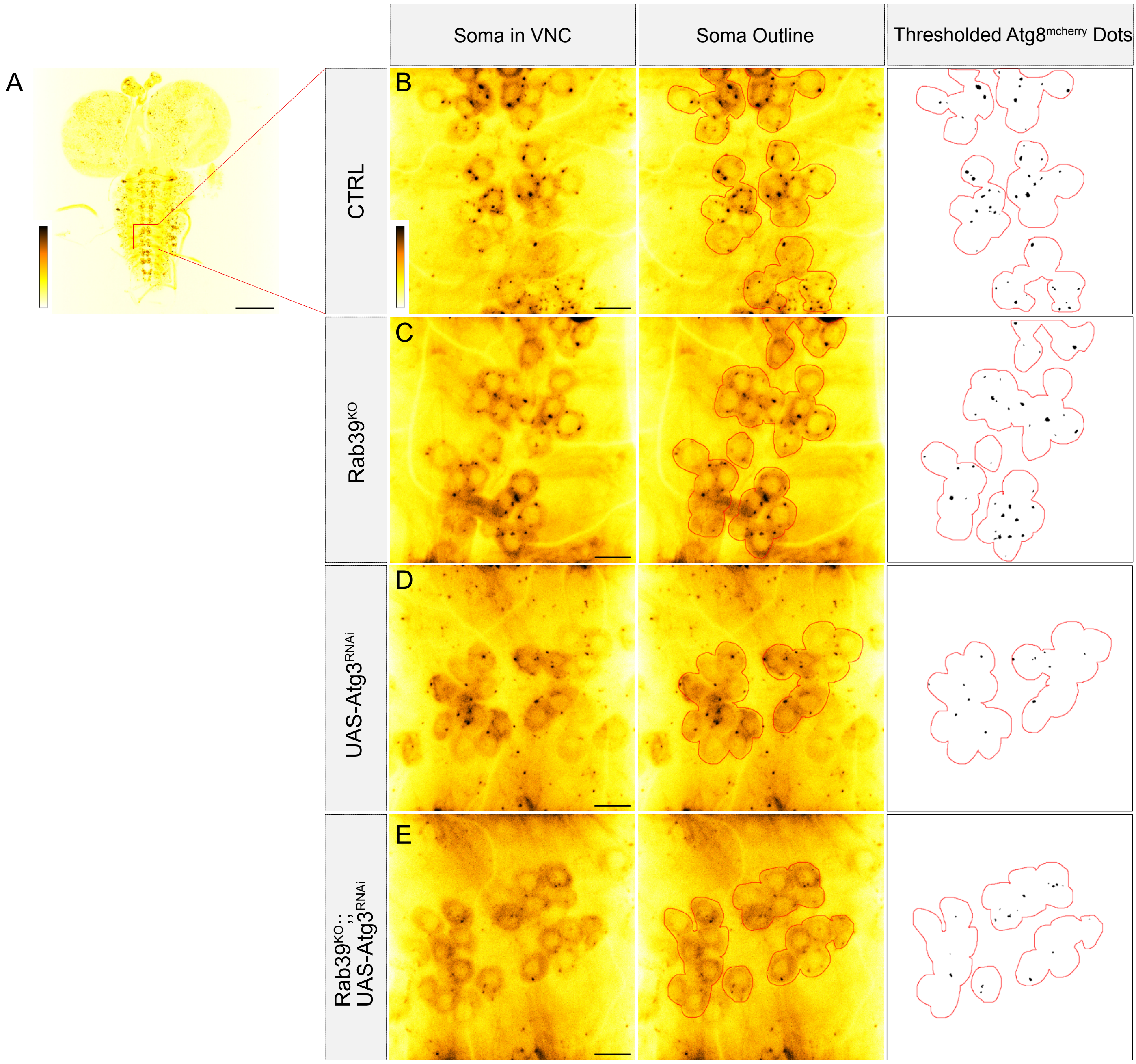


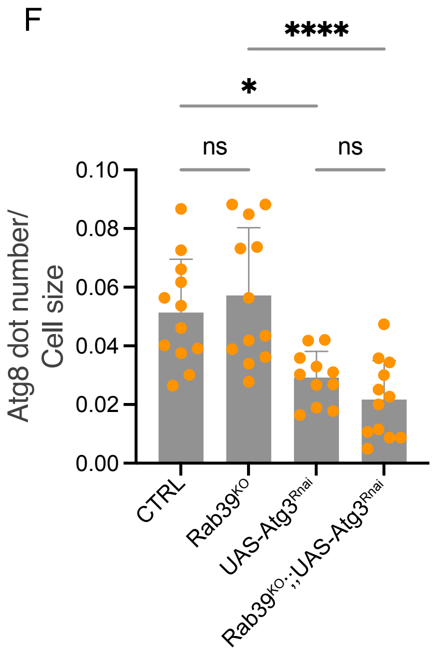

**Figure S3. Synaptic protein levels in control and Rab39$^{KO}$ animals**

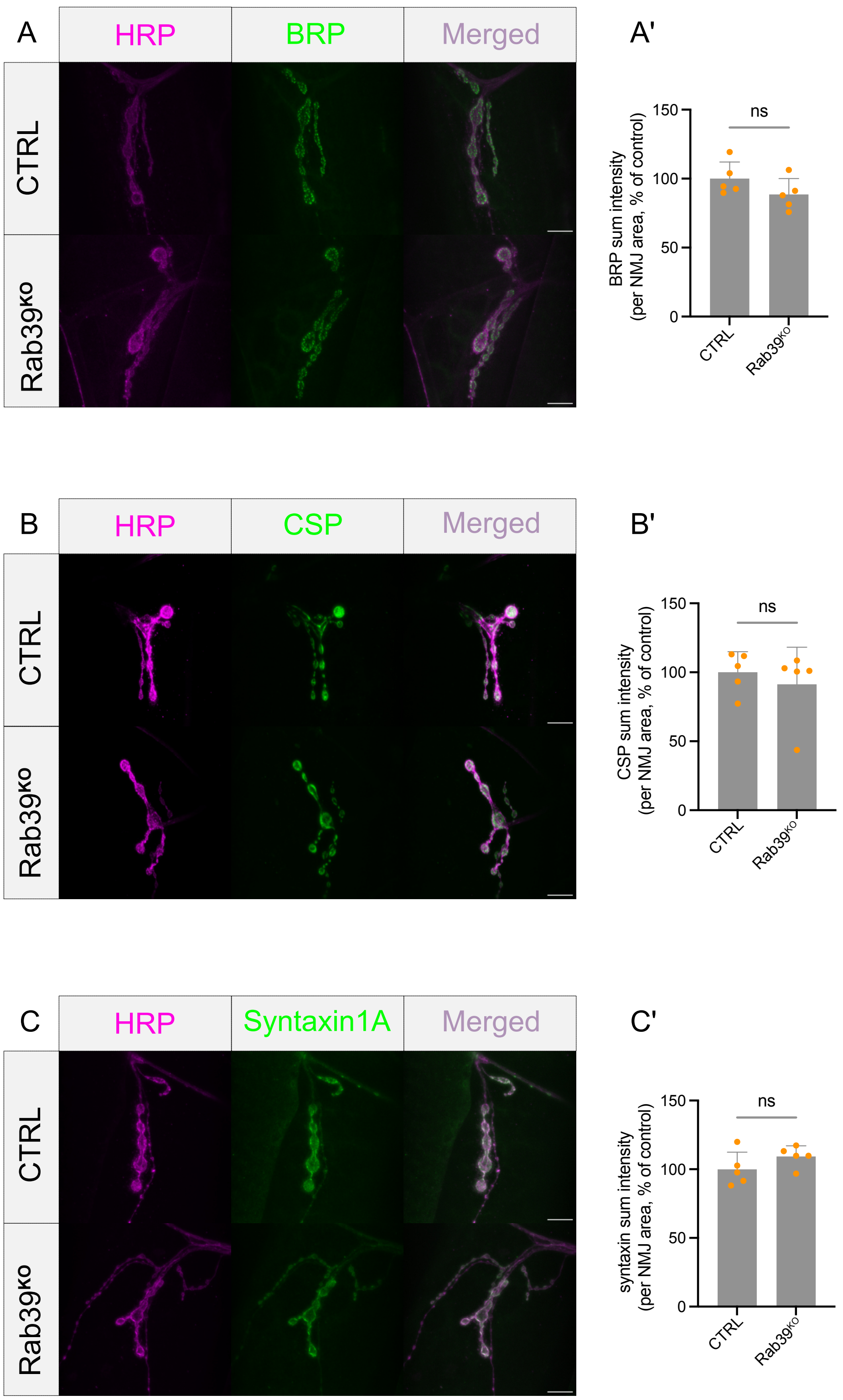

**Figure S4. Pathogenic Rab39B$^{T168K}$ mutant mimics Rab39$^{KO}$ and increases synaptic autophagy**

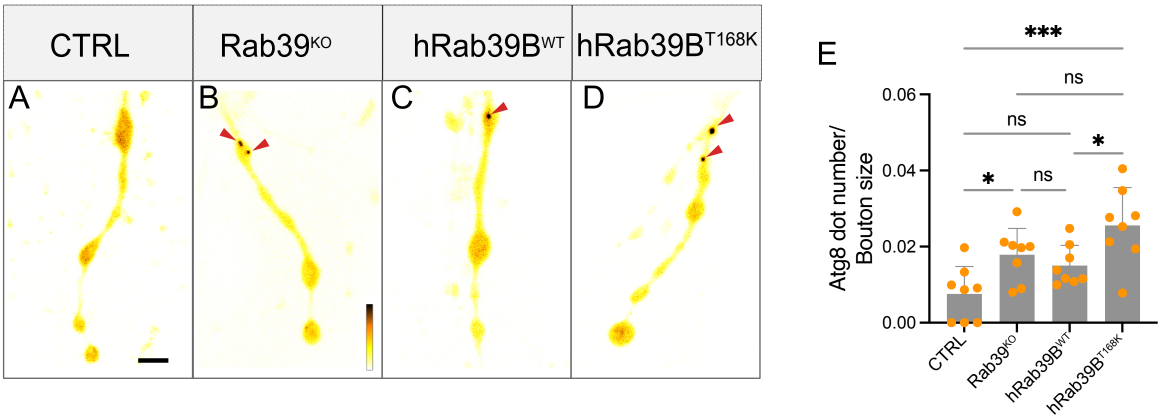

**Figure S5. Localization of Rab39$^{EYFP}$ at the cell bodies of the ventral nerve cord and at NMJs**

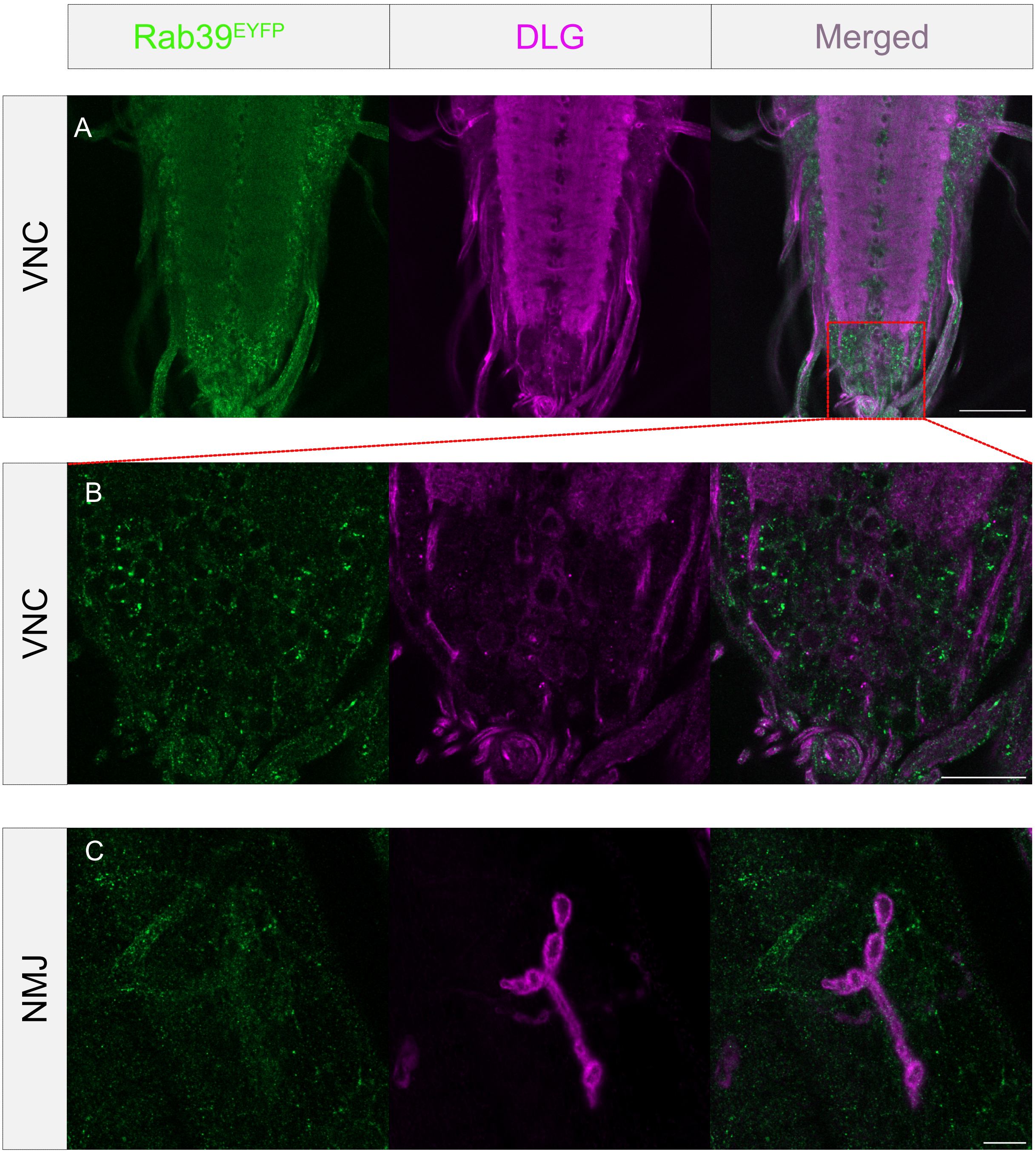
